# Metabolic vulnerabilities in Down syndrome B-cell acute lymphoblastic leukemia can be targeted using Venetoclax

**DOI:** 10.1101/2024.09.30.615872

**Authors:** Dallas Jones, Elena Woods, Zhenhua Li, Kristin L. Schaller, Eric Hoffmeyer, Ben Kooiman, Joseph Fernandez, Tegan Wharton, Dejene Tufa, Spencer C. Hall, George Trahan, Kelly W. Maloney, Holly Pacenta, Kelly Sullivan, Joaquin Espinosa, James R. Roede, John Marentette, Brett Stevens, Anagha Inguva, Daniel Stephenson, Julie A. Reisz, Angelo D’Alessandro, Craig T. Jordan, Jun J. Yang, Michael R. Verneris

## Abstract

Children with Down syndrome (DS) and B-cell acute lymphoblastic leukemia (B-ALL) are at increased risk for treatment-related mortality and relapse, highlighting the need for new therapies. Leukemia cell lines (CLs) have been fundamental to understanding therapeutic responses to pharmacological agents. We generated three DS B-ALL CLs characterized with diverse genomic alterations, including *IGH::CRLF2* rearrangement (*BCR::ABL1 like*), mutations in *FLT3* and *TP53*, and a novel *ERG::CEBPD* rearrangement. DS CLs had diminished proliferation, metabolism, and mitochondrial function when compared to non-DS (NDS) CLs and interestingly, these findings were similar to NDS Philadelphia chromosome-like (Ph-like) B-ALL CLs. Based on similar mitochondrial defects and prior preclinical data using Venetoclax for Ph-like B-ALL, we hypothesized that Venetoclax would be effective in DS. Intriguingly, Venetoclax was more effective in DS when compared to both NDS and Ph-like CLs. Efficacy was observed in DS patient derived xenografts (PDXs) and diagnostic/relapsed patient samples treated with Venetoclax, which synergized with Trametinib and Vincristine. Mass spectrometry-based multiomics analyses in DS and NDS B-ALL patient samples revealed an enriched metabolite profile in DS, particularly in the hubs of glucose metabolism and polyunsaturated phosphatidylcholines and phosphatidylinositols. Transcriptome analyses in DS B-ALL patients (n=249) supported enhanced glucose and fatty acid metabolism. Glucose regulated B-ALL viability through *de novo* serine biosynthesis. Targeting serine synergized with Venetoclax in DS B-ALL CLs. In summary, we have generated novel tools for studying DS B-ALL and identify altered metabolism in DS that responds to Venetoclax.

## Introduction

Down syndrome (DS) is the most common genetic abnormality and is caused by trisomy 21^1^. It is associated with multifaceted clinical and developmental challenges, including upper respiratory complications, congenital heart abnormalities, immunological and hematological disorders, including a 20-fold increased risk of B-cell acute lymphoblastic leukemia (B-ALL)^2^. DS B-ALL patients experience greater treatment related toxicity and relapse leading to inferior survival when compared to non-DS (NDS) B-ALL patients^3-5^. The causes for higher B-ALL prevalence and chemoresistance remain unclear. Surprisingly, there are no DS B-ALL cell lines (CLs) for study. We established three DS B-ALL cell lines to better understand DS biology and explore potential therapeutic interventions.

Recently, we identified that the genomic landscape of DS B-ALL contained 15 distinct subtypes, including a *CCAAT/Enhancer Binding Protein-altered* (*C/EBP-altered*) subtype that co-occurred in >42% of patients with *FLT3* mutation^6^. This novel subtype, and the known *CRLF2-rearranged* (*BCR::ABL1-like*) subgroup, had some of the worst survival rates amongst DS B-ALL patients^6^. The DS B-ALL CLs reported herein represent three diverse subtypes; including 1) a novel *CEBPD* fusion to the chromosome 21 ETS-family transcription factor, *ERG* (*ERG::CEBPD*), that also harbored mutated *FLT3* and *TP53*, 2) *CRLF2-rearranged* (*BCR::ABL1-like*), and 3) a B-other ALL with *TP53* mutation. A hallmark of DS and Philadelphia chromosome-like (Ph-like) B-ALL is the overexpression of CRLF2 (cytokine receptor-like factor 2), found in >50% of DS and <10% of NDS cases, which frequently co-occurs with activating *JAK2* mutations^7-11^. Compared to other subtypes of B-ALL, Ph-like patients also have inferior survival rates^10^.

Recent preclinical therapeutic approaches for Ph-like B-ALL have targeted BCL2 using the BH3-mimetic Venetoclax^12^. Herein, we show similar defects in proliferation and mitochondrial function in DS and Ph-like when compared to NDS B-ALL CLs. Based on these results, we hypothesized that Venetoclax would be active against DS B-ALL. Intriguingly, Venetoclax was more effective in DS, when compared to both Ph-like and NDS CLs. Additionally, Venetoclax sensitivity was observed in DS B-ALL patient samples *in vitro* and in CLs and patient derived xenografts (PDXs) *in vivo*. In contrast to CLs, DS B-ALL patient samples displayed an altered metabolic profile relative to NDS samples that might contribute to their chemoresistance. Finally, we show synergy between Venetoclax and Vincristine at low nanomolar concentrations in DS B-ALL, prompting its consideration into established chemotherapy regiments for DS patients.

## Methods

### Patient samples and cell culture

De-identified NDS and DS B-ALL samples were obtained from bone marrow or peripheral blood samples of patients and families who consented to participate in the University of Colorado leukemia cell bank protocol (COMIRB #4876). NDS (NALM6, HB11;19, KOPN8, RCH-ACV, SEM) and Ph-like B-ALL CLs used (MHHcALL4 and MUTZ5) are all commercially available (ATCC or DSMZ). A detailed list of methods for generating CLs and all drugs used in this study can be found in Supplemental Materials.

### RNA-seq and whole-genome sequencing (WGS)

A full list of details involving RNA-seq and WGS can be found in Supplemental Materials.

### Mouse studies

Xenograft studies were performed in NSG-mice (IACUC 00251) and are detailed in Supplemental Materials. Bioluminescence imaging (BLI) of luciferin treated mice was conducted to monitor leukemia growth of CLs.

### Flow cytometry, Western blot, and Immunoprecipitation

A detailed list of all flow cytometry, western blot, and immunoprecipitation protocols and antibodies used can be found in Supplemental Materials.

### Seahorse Mito-stress test and immunofluorescence imaging

The Seahorse XF cell Mito-stress test kit (Agilent) was used, followed by injection of 2-deoxyglucose. A detailed list of Seahorse and immunofluorescence imaging protocols can be found in Supplemental Materials.

### U-^13^C_6_-glucose tracing and ultra-high-pressure liquid chromatography-mass spectrometry (UHPLC-MS) lipidomics and metabolomics

Analyses were performed as previously described^13^, as detailed in Supplemental Materials.

## Results

### Generation of three DS B-ALL cell lines and their DS subtype alignment

We generated three DS B-ALL CLs (DS-1, DS-76, and DS-81) and performed comprehensive transcriptomic and genomic profiling using RNA-seq and WGS. The three CLs demonstrated distinct gene expression profiles, as visualized using uniform manifold approximation and projection tool (UMAP) in the context of our previously published DS B-ALL dataset^6^ (Figure 1A). DS-1 harbored an *ERG*::*CEBPD* rearrangement, that juxtaposes the downstream of *CEBPD* to exon 3 of *ERG* (Figure 1B), that presumably drives overexpression of *CEBPD*. This CL has a FLT3 mutation (Figure 1C), which is frequently seen in C/EBPalt DS B-ALL^6^. DS-76 has *IGH*::*CRLF2* rearrangement with CRLF2 upregulation, *JAK2* mutation (R683G, Supplemental Figure 1A), and a *BCR*::*ABL1*-like gene expression profile (Figure 1A). No known gene rearrangements were identified in DS-81, and it was classified as B-other. Notably, *TP53* mutations are rare in diagnostic DS B-ALL^6^, and were identified in DS-1 and DS-81, affecting both alleles. DS-1 harbored two TP53 mutations, which are very close to each other and compound heterozygous as they are supported by different reads (Figure 1D). For DS-81, the *TP53* mutation affected one allele with the other allele lost due to 17p deletion (Figure 1D, Supplemental Figure 1B). Full mutational analyses is provided (Supplemental Figure 1A). Next, we confirmed the expression of B-lineage proteins in DS B-ALL CLs compared to commercially available NDS (NALM6 and SEM) and Ph-like B-ALL CLs (MHH-cALL4 and MUTZ5). As expected, all B-ALL CLs expressed CD19, except for DS-1 that lacked surface CD19 and CD22 due to relapse following CD19- and CD19/CD22-CAR T cells (Figure 1E, Supplemental Figure 1C). All DS B-ALL CLs expressed CD79A and CRLF2 expression was confirmed in DS-76, MHH-cALL4, and MUTZ5 (Figure 1E); along with other hematopoietic- and B-lineage proteins (CD45, CD34, CD38, and IL-7Rα, Supplemental Figure 1C). Figure 1F confirms expression of B-lineage transcripts (*CD19, CD79A, EBF1, PAX5, RAG1, RAG2*), *CEBPD*, and *ERG* in DS CLs.

**Figure 1.**
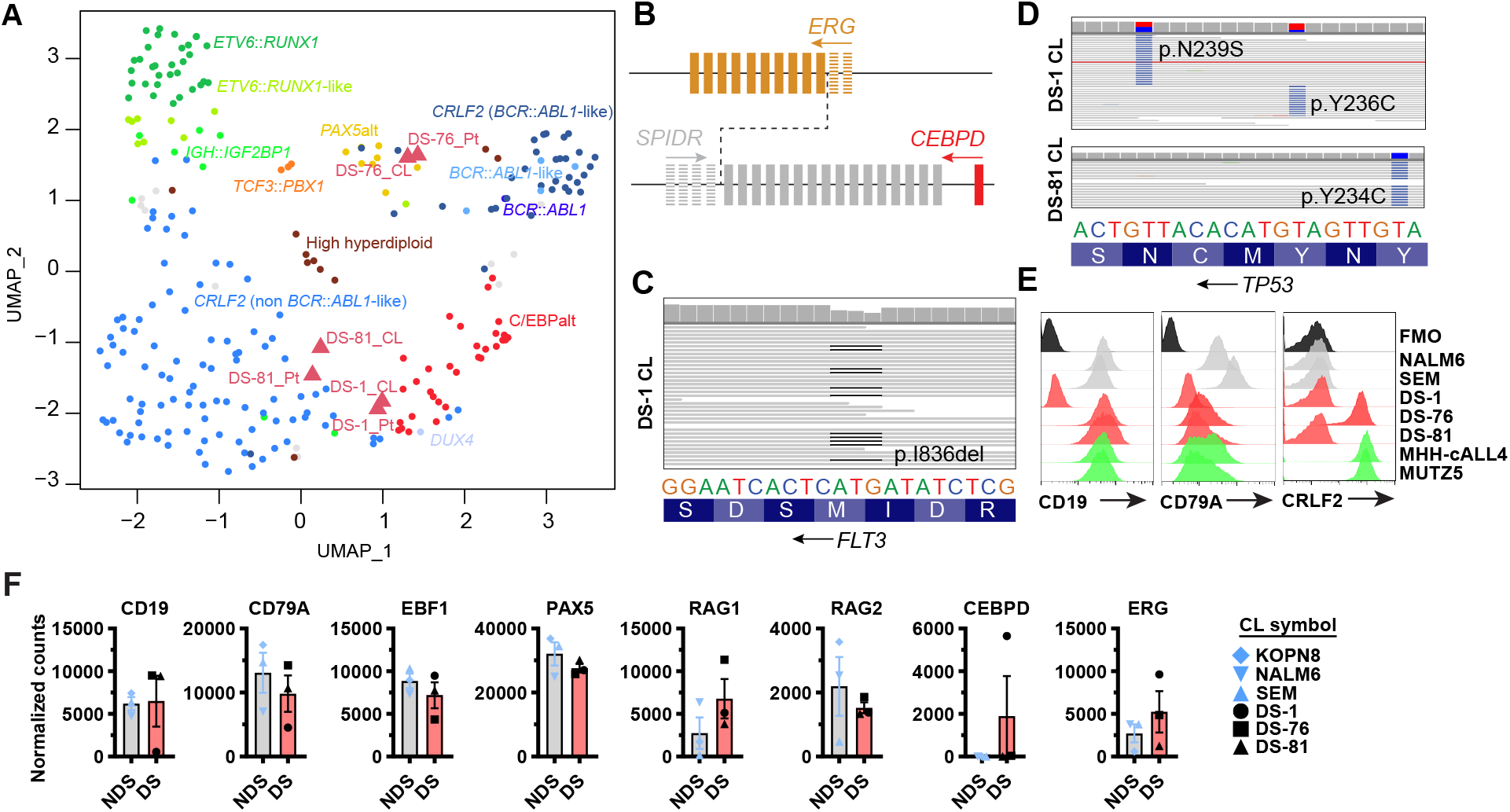
Generation of three DS B-ALL cell lines and their DS subtype alignment. **(A)** Visualization of gene expression profiles of the DS cell lines (CLs) and their corresponding patient samples (Pt) using uniform manifold approximation and projection (UMAP) together with previously published 249 primary DS B-ALL samples. **(B)** ERG::CEBPD rearrangement identified in DS-1. Each box represents an exon, and solid boxes indicate exons involved in the rearrangement. **(C)** FLT3 mutation in DS-1. **(D)** TP53 mutations in DS-1 and DS-81. **(E)** Flow cytometry analyses of surface CD19 and CRLF2 expression and intracellular CD79A expression in NDS, DS, and Ph-like B-ALL CLs. Cells were gated on live singlets. **(F)** RNA-Seq analyses of CD19, CD79A, EBF1, PAX5, RAG1, RAG2, CEBPD, ERG normalized expression levels in NDS (KOPN8, NALM6, and SEM) and DS B-ALL CLs. Bar graphs represent average ± SEM. Wald test and Adj.

### Diminished proliferation and poor mitochondrial function in DS B-ALL cell lines

During the generation of DS B-ALL CLs, we observed delayed growth relative to NDS CLs. DS recapitulated the behavior of the two commercially available Ph-like B-ALL CLs (MHH-cALL4 and MUTZ5), although the small sample size precluded statistical analysis. Nevertheless, cell-trace-violet proliferation dye confirmed that like Ph-like B-ALL CLs, DS CLs had significantly reduced proliferation compared to NDS (Figure 2A). These results were validated *in vivo* using NSG-mice (Supplemental Figure 2A-C). Cell cycle analyses revealed DS had a significantly lower proportion of cells in S-G_2_-M, that coincided with increased proportions in G_1_ when compared to NDS, with Ph-like mimicking DS CLs (Figure 2B). Differential expression of proteins associated with cell signaling and growth (mTOR, P-STAT5, P-ERK1/2, P-AKT, c-Myc) were found amongst the three cohorts that could not explain proliferation defects observed (Supplemental Figure 2D-E). Mutational status of NDS and Ph-like CLs is provided (Supplemental Figure 2F).

**Figure 2.**
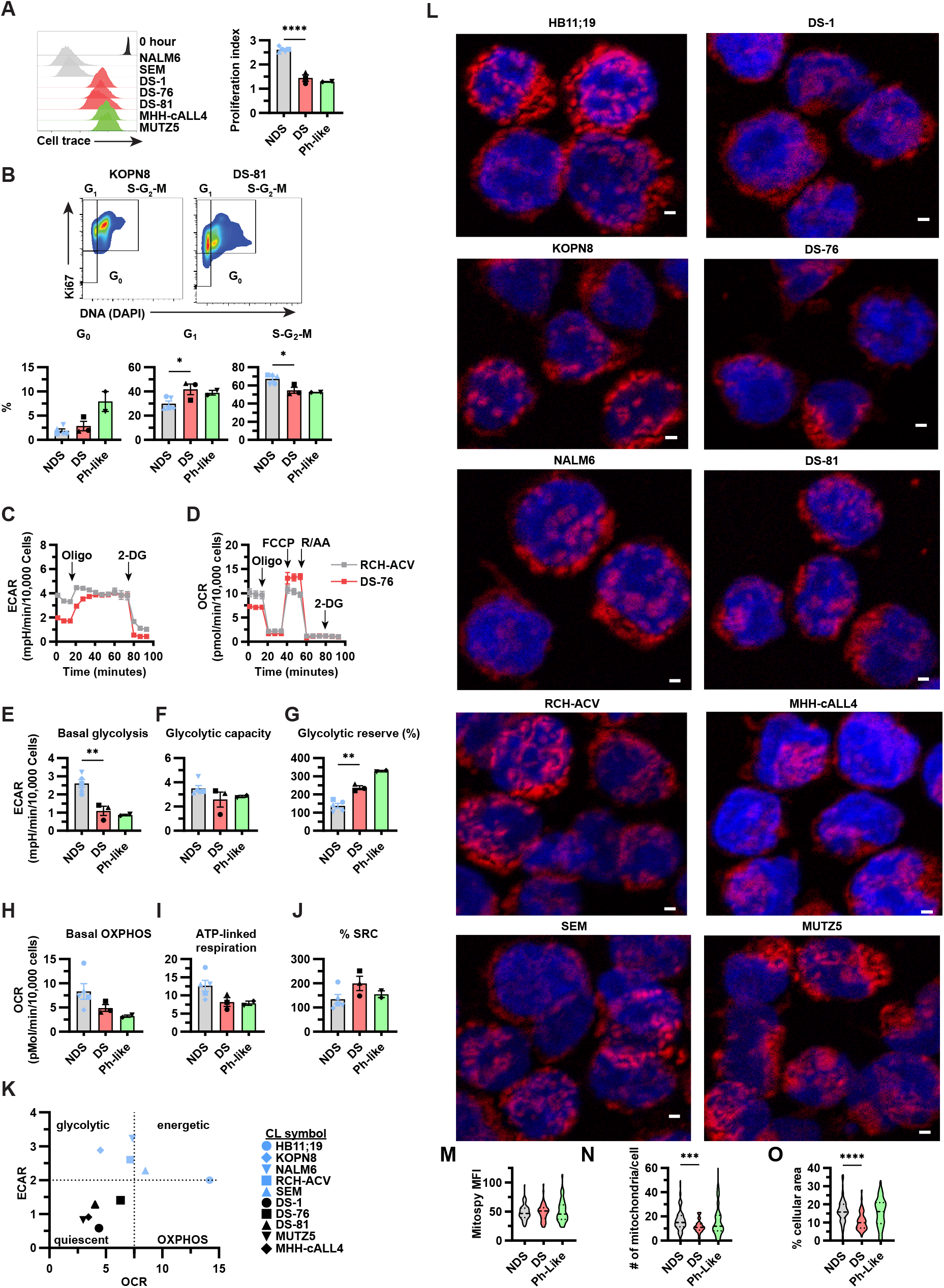
Diminished proliferation and poor mitochondrial function in DS B-ALL cell lines. **(A)** Representative flow cytometry histograms of cell trace violet labeled NDS, DS, and Ph-like B-ALL cell lines (CLs) after 6-days of culture and corresponding graphs summarizing cell proliferation differences. Cells were gated on live singlets. N=3+ replicates from all CLs. **(B)** Representative flow cytometry plots for cell cycle analyses in KOPN8 and DS-81 B-ALL CLs using Ki67 and DAPI expression and corresponding graphs summarizing cell cycle differences. G_0_=Ki67^-^DAPI-single positive; G_1_=Ki67^+^DAPI-single positive; G_2_-S-M=Ki67^+^DAPI-double positive. Cells were gated on live, singlet CD19^+^ cells. DS-1 cells were gated on live singlets. N=3+ replicates from all CLs. **(C-D)** Representative Seahorse Mito-Stress traces for extracellular acidification rate (ECAR; mpH/min/10,000 cells) **(C)** and oxygen consumption rate (OCR; pmol/min/10,000 cells) **(D)**. The timing and addition of mitochondrial inhibitors for ECAR: Oligomycin (Oligo, 6 μM) and 2-deoxyglucose (2-DG 100 mM); along with those for OCR: Oligo (6 μM), FCCP (6 μM), and R/AA (Rotenone 250 nM, antimycin 165 nM) are shown. **(E-G)** ECAR readings for basal glycolysis **(E)**, glycolytic capacity **(F)**, and glycolytic reserve **(G)** from Seahorse experiments. **(H-J)** OCR readings for basal OXPHOS **(H)**, ATP-linked respiration **(I)**, and % spare respiratory capacity (SRC) **(J)** from Seahorse experiments. N=5+ replicates from all CLs. **(K)** Seahorse energy phenotype in CLs using basal glycolysis and basal OXHPOS. **(L-O)** Confocal microscopy for mitochondrial morphology using Mitospy (red) and nuclear DNA using DAPI (blue) **(L)**. Images are maximum intensity z-stack projections after background subtraction (20 pixel rolling ball radius), and enhanced local contrast (blocksize 9, slope 4) (FIJI). Results representative of n=17-75 cells from each CL. **(M-O)** Violon plots for Mitospy mean fluorescence intensity (MFI) expression levels **(M)**, # of mitochondria/cell **(N)**, and % cellular area **(O)** in NDS, DS, and Ph-like B-ALL CLs from **L**. All bar graphs represent average ± SEM. Statistical significance was determined using two-tailed T-test **A, B, E-J** and Mann-Whitney U test **M-O**. *p<0.05, **p<0.01, ***p<0.001, and ****p<0.0001.

Next we assessed cellular bioenergetics using Seahorse mitochondrial-stress assay modified to include a 2-deoxyglucose injection (Figure 2C-D). A profound impact on glycolysis was observed in DS B-ALL CLs as basal glycolysis was significantly reduced, glycolytic capacity unaffected, and glycolytic reserve significantly elevated when compared to NDS (Figure 2E-G), with Ph-like following the same trends as DS. Moreover, basal oxidative phosphorylation (OXPHOS) and ATP-linked respiration were diminished in DS and Ph-like CLs, whereas spare respiratory capacity (SRC) was slightly elevated when compared to NDS (Figure 2H-J). Overall, Seahorse experiments identified a quiescent phenotype in DS and Ph-like CLs, while NDS appeared glycolytic and energetic (Figure 2K). Confocal microscopy confirmed a decreased mitochondrial network in DS compared to NDS (Figure 2L-O). Oxidative stress was diminished in DS and Ph-like CLs compared to NDS (Supplemental Figure 2G). Interestingly, RNA-seq comparisons between DS and NDS B-ALL CLs showed only 6 chromosome 21 genes upregulated in DS (Supplemental Figure 2H). Notably, Ingenuity pathway analysis (IPA) showed mitochondrial dysfunction as the top canonical pathway altered in DS (Supplemental Figure 2I). Collectively, these data show decreased proliferation in DS CLs and an impairment of both glycolytic metabolism and mitochondrial function promoting a state of metabolic quiescence.

### Altered glucose metabolism in DS B-ALL cell lines

Since Seahorse mitochondrial-stress assays and observations of cells in culture (Figure 3A) suggested declines in basal glycolysis in DS and Ph-like B-ALL CLs compared to NDS, we further interrogated glucose metabolism using stable isotope labeled glucose tracing by mass spectrometry. This allowed us to identify glucose flux through glycolysis (i.e. generation of lactate as a glycolytic endpoint or as pyruvate for further metabolism in the tricarboxylic acid cycle (TCA)) versus the ancillary pentose phosphate pathway (PPP) for NADPH generation and redox balance. Using partial least-squares discriminant analysis (PLS-DA) we observed distinct differences in glucose metabolism amongst DS, Ph-like, and NDS CLs (Figure 3B). Total levels of intracellular glucose were significantly higher in DS B-ALL than in the other CLs tested (Figure 3C). As expected, DS had significant decreases in total lactate when compared to NDS, while total pyruvate levels were similar between the three cohorts (Figure 3C). Glucose tracing also revealed that Ph-like B-ALL cells direct glucose into the PPP as its activation, indicated by the labeled levels of [^13^C_6_]phosphogluconate normalized to precursor [^13^C_6_]hexose phosphate, was higher in the MUTZ5 CL when compared to DS and NDS (Figure 3C). These results have been previously reported from network analyses in Ph-like B-ALL^12^. A full summary of glucose tracing experiments revealed downregulation of most glycolytic and TCA intermediates in DS and Ph-like B-ALL CLs (Supplemental Figure 3, 4); but minimal alterations in PPP aside from its activation in Ph-like B-ALL (Supplemental Figure 5). This confirmed that glucose metabolism via glycolysis and subsequent entry into the TCA cycle is impaired in DS B-ALL CLs, but highly active in NDS.

**Figure 3.**
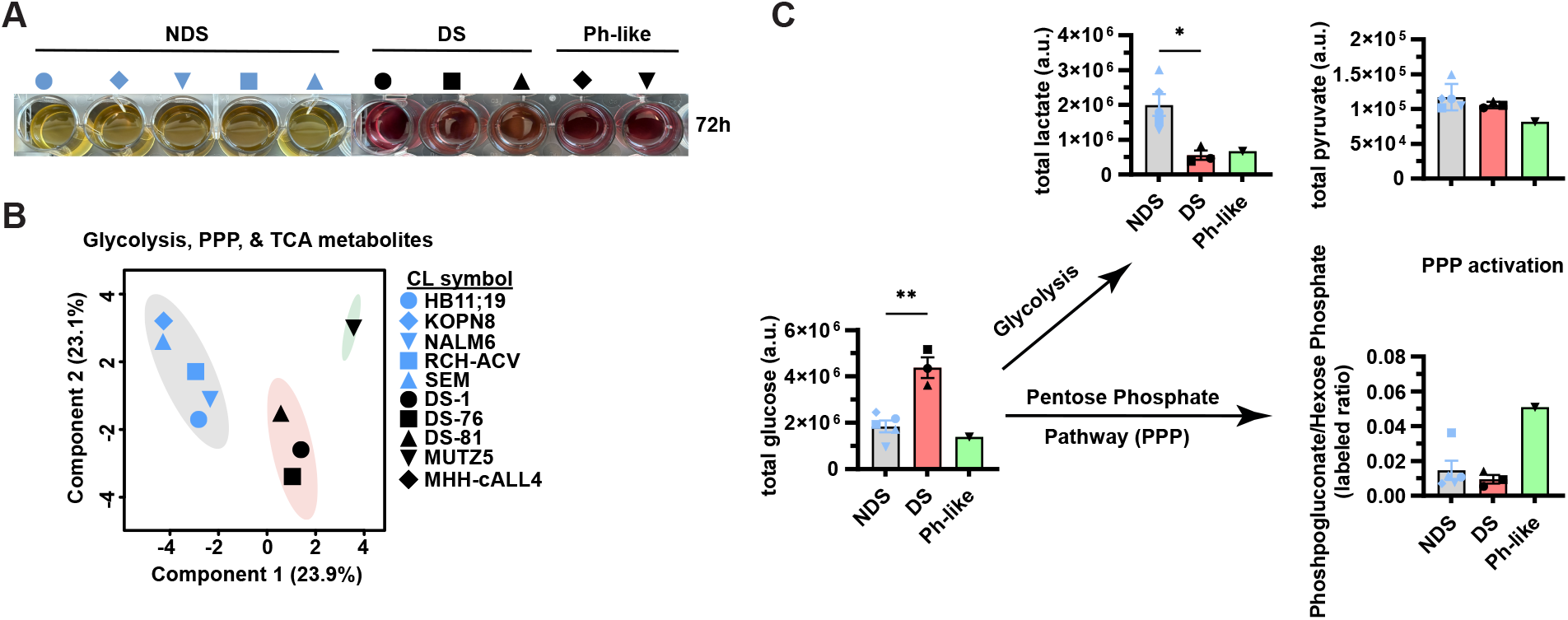
Altered glucose metabolism in DS B-ALL cell lines. **(A)** Images of cell culture media changes in NDS, DS, and Ph-like B-ALL cell lines (CLs) cultured for 72h at 1 ×10^6^ cells/ml. **(B)** NDS, DS, and MUTZ5 Ph-like B-ALL CLs were cultured with [^13^C_6_]glucose for 24h before cells were analyzed by mass spectrometry. Changes in metabolic phenotypes among groups are presented via PLS-DA (partial least squares-discriminant analysis). **(C)** Summary of glucose tracing experiments from **B** depicting total levels of glucose, along with its differential usage for glycolysis (total lactate and total pyruvate) or pentose phosphate pathway (PPP) based on the [^13^C_6_]phosphogluco-nate/[^13^C_6_]hexose phosphate ratio as a readout for PPP activation. Data is presented in arbitrary units (a.u.). All CLs were analyzed in triplicate and bar graphs represent average ± SEM. Statistical significance was determined using two-tailed T-test. *p<0.05 and **p<0.01.

### Altered metabolism in DS B-ALL patients

Since DS B-ALL CLs demonstrated a unique disruption to glycolytic and mitochondrial metabolism, we assessed the impact of adaptation to cell culture on metabolic state and cell cycle progression using patient samples from DS and NDS cohorts. No cell cycle differences were found in these patient samples (Figure 4A). Conversely, we found CLs proliferate at significantly higher rates than patient samples as <16% of patient samples were in S-G_2_-M, whereas CLs were >50% (Figure 4A). Supplemental Figure 6 provides a list of patient features. As expected, CD19^HI^ B-ALL patient cells (top 1% and 10%) had significantly higher proliferation rates than bulk CD19^+^ cells (Supplemental Figure 7A), owing to its role in B-ALL proliferation^14^. These results were the same for CRLF2-rearranged patients as CRLF2^HI^ cells were also CD19^HI^. Using these same comparisons, we show similar proliferation rates in DS B-ALL CLs as their matching patients CD19^HI-1%^ cells (or CD34^HI-1%^ cells for DS-1, Supplemental Figure 7B), consistent with clonal selection during CL generation, that was also observed when comparing matching PDXs and patient samples (DS-2 and DS-3, Supplemental Figure 7C). Cryopreservation had no effect on cell cycle in CLs (Supplemental Figure 7D).

**Figure 4.**
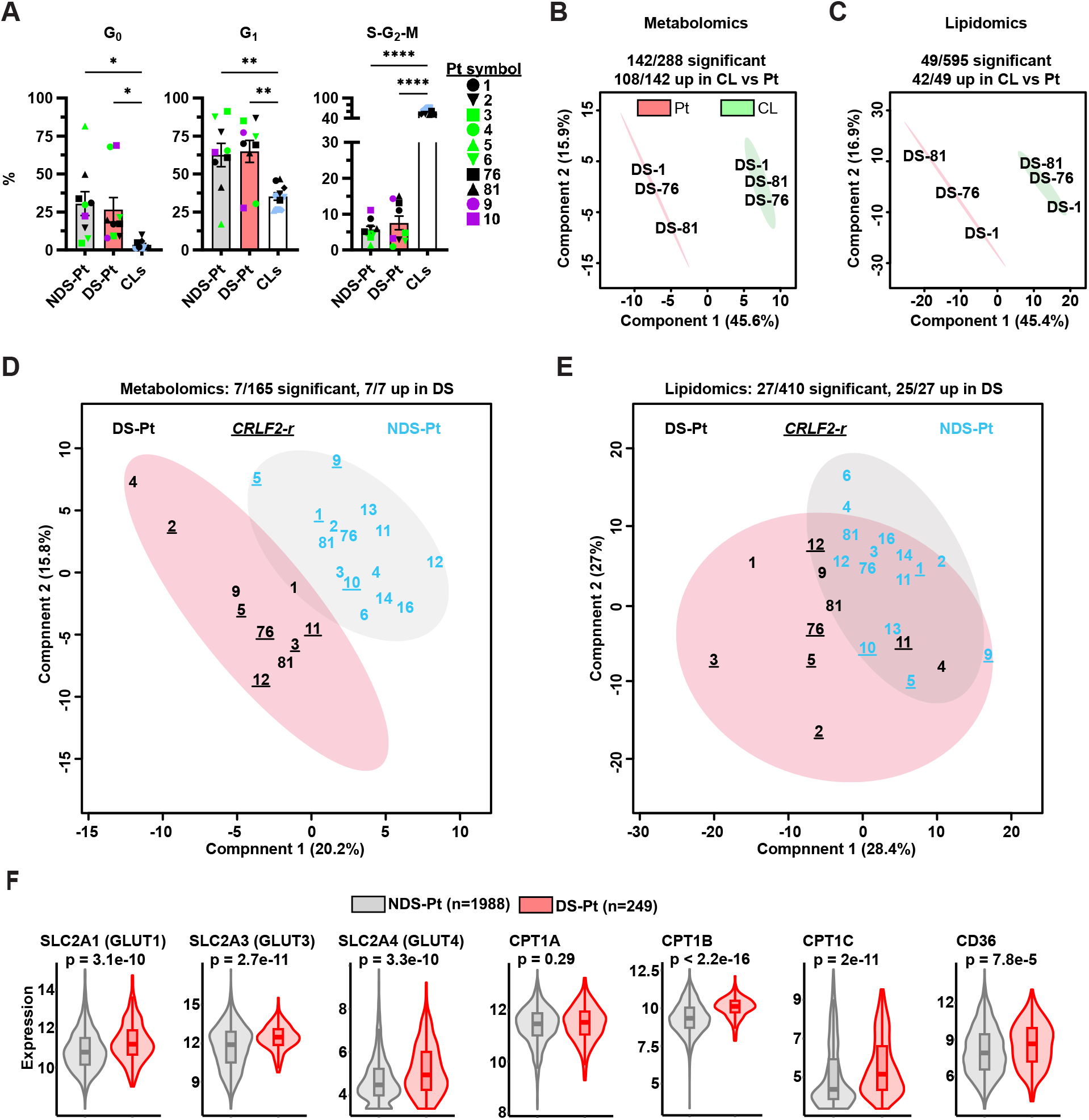
Altered metabolism in DS B-ALL patients. **(A)** Summary of cell cycle analyses in B-ALL cell lines (CLs) and NDS and DS B-ALL patient (Pt) samples. CLs were gated on live singlet CD19^+^ cells (except for DS-1 gated on live singlets). Patient samples were gated on live, singlet, CD34^+/-^, CD45^+/-^, CD3^-^, CD66B^-^, CD14^-^, CD19^+^. CRLF2^+^ cells were defined as > 80%-positive. DS-1 patient sample was gated on CD45^+/-^, CD3^-^, CD66B^-^, CD14^-^, CD19^-^, CD34^HI^ as this represented >80% of cells. Cells were further stained with Ki67 and DAPI. G0=Ki67^-^DAPI^-^single positive; G1=Ki67^+^DAPI-single positive; G2-S-M=Ki67^+^DAPI-double positive. **(B-C)** Mass spectrometry was performed on matching DS B-ALL CLs (green) and Pt samples (red), then data subjected to PLS-DA (partial least squares-discriminant analysis) for differences in molecular composition of metabolites **(B)** and lipids **(C)**. Patient samples were thawed from cryopreservation, flow cytometry stained, and sorted as described in **A. (D-E)** Mass spectrometry was performed on flow cytometry sorted NDS and DS B-ALL patient samples as described in **A**, then data subjected to PLS-DA for differences in molecular composition of metabolites **(D)** and lipids **(E)**. Numbers denote unique patients with NDS-Pt=blue numbers and DS-Pt=black numbers. *<u>CRLF2-r</u>* patient numbers are underlined. **(F)** Violin plots for the expression levels of SLC2A and CPT1 family members along with CD36 using RNA-seq from NDS (n=1988) and DS (n=249) B-ALL patients. All bar graphs represent average ± SEM. Statistical significance was determined using one-way ANOVA with Tukey posttest in **A**, two-tailed T-test in **B-E**, and Wilcoxon rank-sum test in **F**. *p<0.05, **p<0.01, ***p<0.001, and ****p<0.0001.

Next, we used mass spectrometry-based metabolomics and lipidomics in matching DS B-ALL CLs and flow cytometry sorted patient samples. PLS-DA of metabolomic data revealed distinct differences between CLs and patient samples, as 142/288 metabolites were significantly altered, with 108/142 elevated in CLs (Figure 4B). PLS-DA of lipidomic data showed a similar pattern, as 49/595 lipids were significantly altered, with 42/49 elevated in CLs versus patients (Figure 4C). These results corroborate the above proliferation findings, as most of the elevated lipids in CLs were involved in cell membrane structure required in rapidly dividing cells. A list of metabolites and lipids are provided (Supplemental Figures 8, 9).

To examine metabolic differences in DS and NDS B-ALL patient samples, we used this same mass spectrometry approach as it provides a “snapshot” of *in vivo* metabolism. Our only criteria for the NDS B-ALL patient cohort was maintenance of 46 chromosomes. Flow cytometry sorted cryopreserved patient samples from both cohorts (>97% purity) were used. PLS-DA of metabolomic analysis showed elevated levels of metabolites in trisomy 21 as 7/165 metabolites found were significantly changed with all seven being upregulated in DS (Figure 4D). These seven metabolites represent distinct metabolic pathways involved in glucose metabolism (glycolysis, PPP, and amino sugars), glutathione homeostasis, Arg/Pro synthesis, Indole/Trp metabolism, and nucleotides (Supplemental Figure 10A). The PLS-DA of lipidomic data identified greater lipid content in trisomy 21 as 27/410 lipids detected were significantly changed with 25/27 increased in DS patients, including multiple polyunsaturated phosphatidylcholines and phosphatidylinositols (Figure 4E). A list of metabolites and lipids is provided (Supplemental Figure 10B, 11). Next, we evaluated glucose and fatty acid metabolism gene expression from our diagnostic DS B-ALL patient cohort (n=249)^6^ in comparison to diagnostic NDS patients (n=1988)^15^. Transcripts from DS B-ALL patients support an altered metabolic phenotype as genes involved in glucose uptake (SLC2A1, SLC2A3, and SLC2A4; GLUT family of glucose transporters), fatty acid oxidation (CPT1B and CPT1C), and fatty acid uptake (CD36) were significantly upregulated (Figure 4F). Overexpression of GLUT1 was confirmed in DS B-ALL patients using flow cytometry (Supplemental Figure 12A). CD36, GLUT, and CPT1 family member expression levels were similar amongst the three CL cohorts (Supplemental Figure 12B-E). Collectively, we identified an altered metabolic phenotype in DS B-ALL patient samples, comprising glucose metabolism, lipids, and fatty acids.

### Metabolic vulnerabilities in DS B-ALL can be therapeutically exploited using Venetoclax

Based on promising preclinical data utilizing Venetoclax for Ph-like B-ALL^12^ and similar alterations in mitochondrial function between Ph-like and DS B-ALL CLs, we hypothesized that Venetoclax would be effective in DS. We found low IC_50_ values for Venetoclax and Vincristine (<0.04 μM, Supplemental Figure 13A) in DS CLs and compared equal doses of these drugs in the three CL cohorts. The MEK-1/2 inhibitor, Trametinib, that was recently found active in DS B-ALL^16^ had a substantially higher IC_50_ in DS CLs (>1μM, Supplemental Figure 13A). We compared the efficacy of these drugs along with Dexamethasone, Methotrexate, and VU0661013 (targets MCL1) in B-ALL CLs. Intriguingly, only Venetoclax was significantly more effective in DS B-ALL compared to NDS (Figure 5A and Supplemental Figure 13B), with Ph-like CLs showing minimal response to Venetoclax, that matched previous IC_50_ values in these^12^ and NDS^17^ CLs tested. These results were recapitulated *in vivo* as BLI showed diminished leukemia after 4-weeks of Venetoclax treatment (100 mg/kg/day, oral gavage, 5 days/week; Figure 5B) and improved survival in all three DS B-ALL groups (Figure 5C). A summary of BLI over the treatment period is provided (Supplemental Figure 13C).

**Figure 5.**
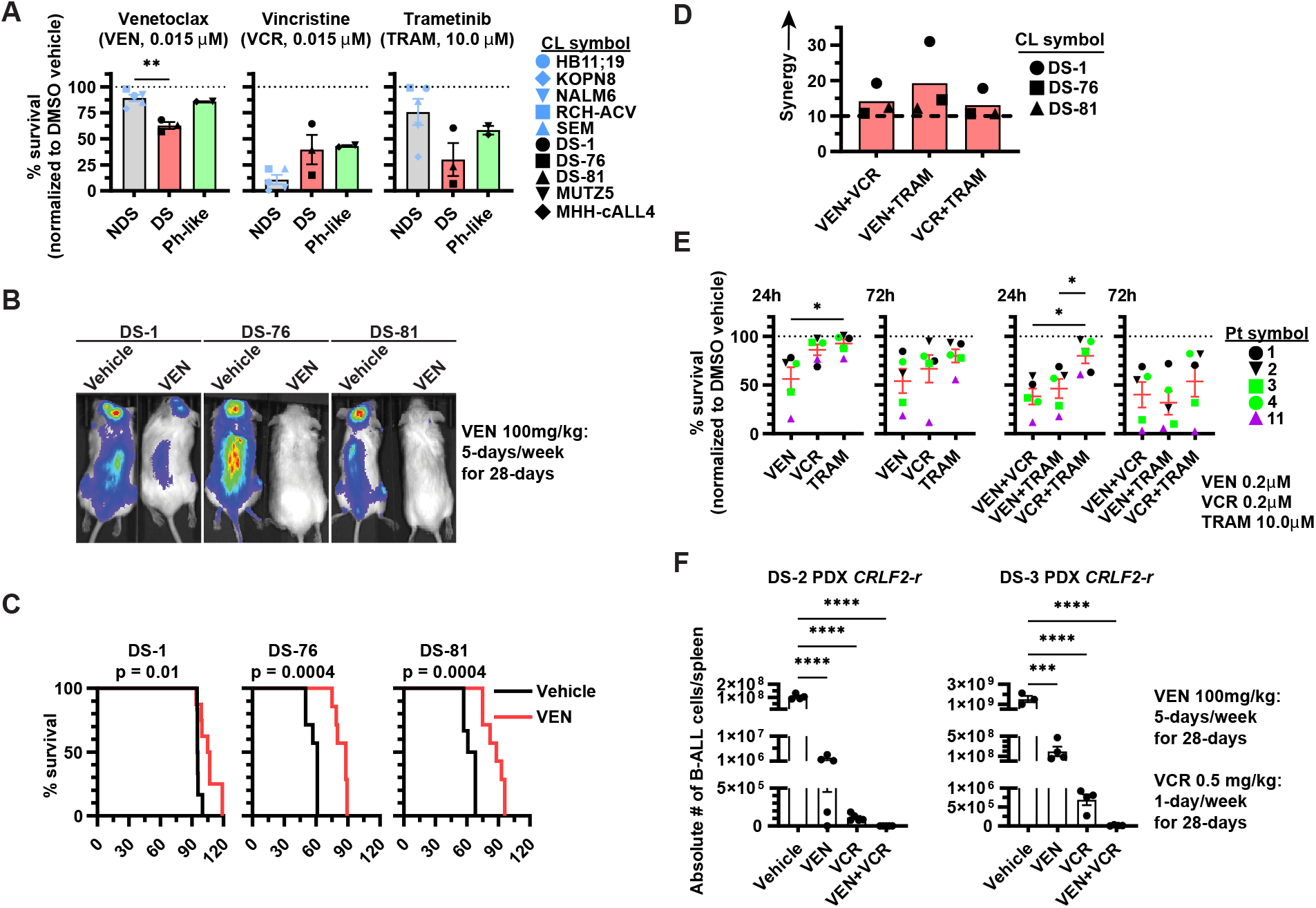
Metabolic vulnerabilities in DS B-ALL can be therapeutically exploited using Venetoclax. **(A)** Graphs for % survival in response to Venetoclax (VEN 0.015 μM), Vincristine (VCR 0.015 μM), and Trametinib (TRAM 10.0 μM) in NDS, DS, and Ph-like B-ALL CLs treated for 72h. Cells were stained with AO/PI dye (acridine orange/propidium iodide) and viability determined using a Cellaca automated cell counter. % survival=viable drug/viable vehicle. Bar graphs represent average % survival ± SEM from n=3 replicates for all CLs and treatments. **(B-C)** NSG mice were tail-vein injected with 1.0 × 10^6^ DS B-ALL CLs from DS-1, DS-76, or DS-81 expressing GFP-Luciferase lentivirus. Then, 2-weeks after injection, treatment with vehicle control or Venetoclax (100 mg/kg/day, oral gavage, 5 days/week) was administered for 4-weeks. **(B)** Representative BLI after 4-weeks of Vehicle or VEN treatment in DS-1, DS-76, or DS-81 mice. **(C)** Kaplan-Meyer survival curve from **B** comparing Vehicle (black) and VEN (red) treatment in DS-1, DS-76, or DS-81 NSG mice. 6-8 mice were used for all CLs and treatment groups. **(D)** Summary of synergy scores for combinatorial drug treatments using VEN, VCR, and TRAM from <u>synergyfinder.fimm.fi</u> in DS B-ALL CLs treated for 72h. All CLs were treated with their IC_50_ values and a 4-fold increase/decrease in these values. All treatments were conducted in triplicate. **(E)** DS B-ALL patient samples were thawed and rested over night before treatment with single dose or combinatorial dose of VEN (0.2 μM), VCR (0.2 μM), or TRAM (10 μM) for 24-or 72h. All cells in the FSCa/SSCa gate were analyzed for singlet, CD34^+/-^, CD45^+/-^, CD3^-^, CD66B^-^, CD14^-^, CD19^+^. CRLF2^+^ cells were defined as > 80%-positive. DS-1 patient sample was gated on CD45^+/-^, CD3^-^, CD66B^-^, CD14^-^, CD19^-^, CD34^HI^ as this represented >80% cells. Cells were stained with annexin-V and DAPI to monitor cell death. Live cells=annexin-V^-^/DAPI^-^. Red lines represent average % survival ± SEM. **(F)** DS-2 and DS-3 PDXs were tail-vein injected into NSG mice as in **B-C**. Flow cytometry was used to assess B-ALL burden in peripheral blood, when levels of human CD19^+^ reached >1% treatment with VEN as in **B-C**, VCR 0.5 mg/kg/week, VEN+VCR combo, or Vehicle treatment began (4-weeks). On day-31 of treatment, mice were sacrificed and B-ALL burden in spleens determined using flow cytometry and cell count beads (absolute # of B-ALL cells/spleen). Statistical significance was determined using a two-tailed T-test in **A**; Log rank test in **C**; and one-way ANOVA with Tukey posttest in **E-F**. *p<0.05, **p<0.01, and ****p<0.0001.

Given the synergy reported between Vincristine and Trametinib in DS PDXs^16^, we tested Venetoclax, Vincristine, and Trametinib in our DS B-ALL CLs and found synergy with all three drugs tested in two-drug combinations (Figure 5D). Venetoclax was effective in DS B-ALL patient samples and displayed similar synergy (Figure 5E). Next, we tested the efficacy of Venetoclax and Vincristine (0.5mg/kg/week) *in vivo* for 4-weeks using 2 DS B-ALL PDXs (DS-2 and DS-3, both *CRLF2-rearranged*). Treatment began once B-ALL was >1% in peripheral blood and on day-31 mice were sacrificed and the absolute # of B-ALL cells/spleen determined. We observed significant reductions in B-ALL with all drug treatments compared to vehicle in both PDXs (Figure 5F). The combination of Venetoclax and Vincristine resulted in >10^5^-fold reduction in splenic B-ALL cells compared to vehicle. Overall, Venetoclax is active against DS B-ALL and synergistic with both Vincristine and Trametinib.

### Differential BCL2 family member expression patterns between cell lines and patient samples

Regulation of mitochondrial apoptosis is mediated by the ratio of anti-apoptotic (BCL2, MCL1, BCL-XL, BAG1) and pro-apoptotic (BAX, BAK, BAD, BIM, NOXA, PUMA) BCL2 family members^18^. Alterations in these might help explain the Venetoclax sensitivity in DS B-ALL. Western blot revealed no alterations in any BCL2 family members between DS and NDS B-ALL CLs, including P-BCL2^SER-70^, which enhances its anti-apoptotic function^19,20^ (Figure 6A-B). Immunoprecipitation experiments in Venetoclax treated DS and NDS CLs showed BCL2 bound strictly to BIM (Supplemental Figure 14).

**Figure 6.**
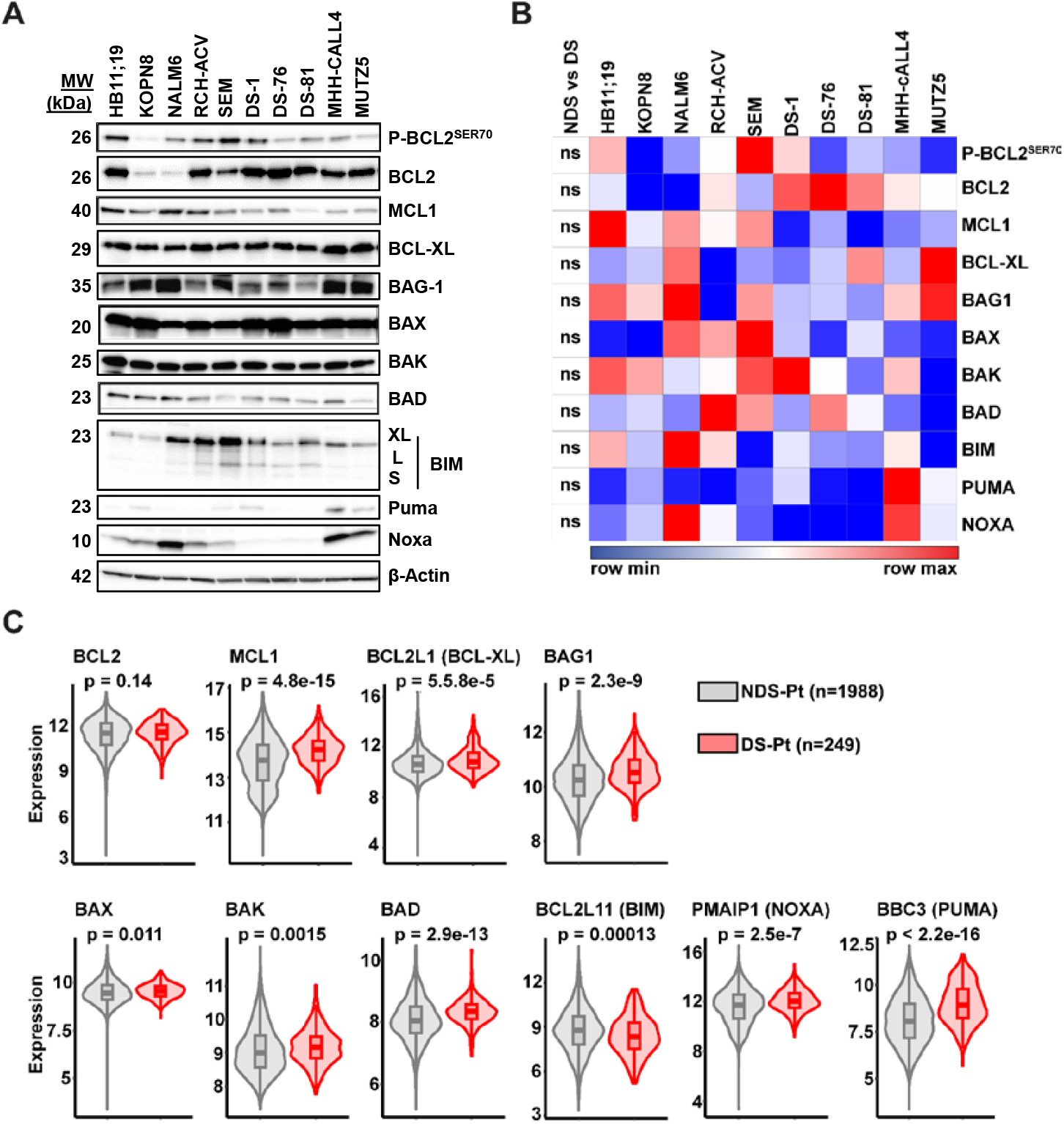
Differential BCL2 family member expression patterns between cell lines and patient samples. **(A)** Representative western blot analyses of Phospho-(P) BCL2 (SER-70), BCL2, MCL1, BCL-XL, BAG1, BIM (XL-, L-, and S-isoforms), BAX, BAK, BAD, PUMA, and NOXA expression levels in whole cell lysates from NDS, DS, and Ph-like B-ALL cell lines. β-Actin was used as a loading control. Western blots were conducted using an n=2+ replicates for each protein analyzed. **(B)** Summary of western blot analyses for BCL2 family member proteins from **A**. Heatmaps represent normalized BCL2 family member expression levels from Image J analyses. Statistical significance was determined using a two-tailed T-test. **(C)** Analyses of BCL2 family member genes from **A** using RNA-seq comparing expression levels in DS (n=249) and NDS (n=1988) B-ALL patients. Wilcoxon rank-sum test used for statistics.

Next, we evaluated BCL2 family member gene expression in DS B-ALL patients and found that *BCL2* expression was not altered between DS and NDS patients; however, *MCL1, BCL2L1* (BCL-XL), and *BAG1* expression levels were significantly elevated in DS (Figure 6C). Pro-apoptotic *BAX, BAK, BAD, PMAIP1* (NOXA), and *BBC3* (PUMA) were significantly upregulated in DS patients, whereas *BCL2L11* (BIM) was downregulated (Figure 6C). Due to batch differences in RNA-seq library prep we did not make comparisons with our DS B-ALL CLs or patient samples. Collectively, differences in BCL2 family members cannot explain the enhanced sensitivity to Venetoclax observed in DS B-ALL CLs; however, DS patients appear reliant on BCL2 family members to resist apoptosis as 3/4 anti-apoptotic and 5/6 pro-apoptotic genes were upregulated relative to NDS.

### *De novo* serine biosynthesis from glucose is critical for B-ALL viability and its blockade synergizes with Venetoclax in DS B-ALL cell lines

We have previously shown Venetoclax targets OXHPHOS-dependent leukemic stem cells in acute myeloid leukemia (AML)^21^; therefore, we next assessed the impact of Venetoclax on DS B-ALL using Seahorse Mito-stress test. Venetoclax had profound effects on DS B-ALL CLs as glycolytic capacity, glycolytic reserve, basal OXPHOS, and ATP were all significantly downregulated (Figure 7A). In contrast, Venetoclax had minimal impact on any Seahorse parameters in NDS and Ph-like B-ALL CLs aside from elevating basal glycolysis in NDS (Figure 7A). Since glucose and lipids were elevated in DS B-ALL patients, we next examined whether manipulation of these pathways mediated survival in B-ALL. Figure 7B depicts the experimental design. Strikingly, glucose was required to maintain viability in all CLs (Figure 7C) and similar IC_50_ values for 2-DG and DRB18 were found (Supplemental Figure 15A). Figure 7D shows the various metabolic fates of glucose, which is required for *de novo* serine biosynthesis as inhibiting only this pathway significantly reduced B-ALL survival in all 3 CL cohorts (Figure 7E). Targeting serine was the only glucose metabolic pathway to enhance Venetoclax efficacy (Figure 7F). CRISPR of phosphoglycerate dehydrogenase (PHDGH; enzyme for serine biosynthesis) in DS B-ALL CLs recapitulated these effects (Figure 7G-H). Glucose tracing confirmed *de novo* serine biosynthesis in B-ALL CLs (Supplemental Figure 15B). The effects of CRISPR on DS B-ALL viability are provided (Supplemental Figure 15C). In summary, Venetoclax impacts glycolytic metabolism in DS B-ALL that is mediated through serine, while all 3 cohorts of CLs required glucose and *de novo* serine biosynthesis for survival.

**Figure 7.**
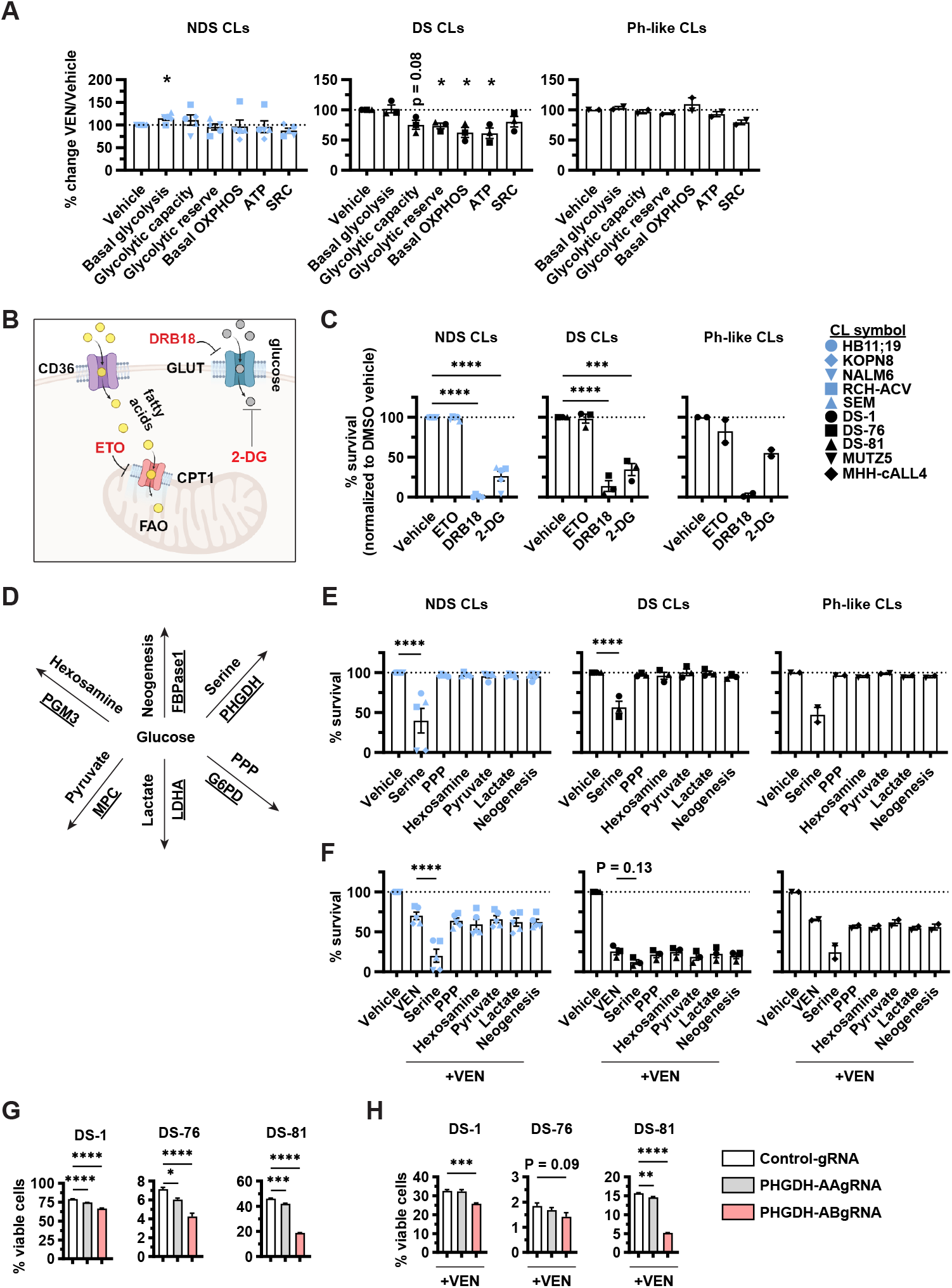
*De novo* Serine biosynthesis from glucose is critical for B-ALL viability and its blockade synergizes with Venetoclax in DS B-ALL cell lines. **(A)** Summary of Seahorse Mito-stress test results after 4h treatment with Venetoclax (VEN). Graphs represent % change VEN/Vehicle for basal glycolysis, glycolytic capacity, glycolytic reserve, basal OXPHOS, ATP, and spare respiratory capacity (SRC) in NDS, DS, and Ph-like B-ALL cell lines (CLs). CLs were treated with VEN 0.02 μM. **(B)** Diagram depicting glucose uptake via the GLUT receptors and its blockade using DRB18 that targets GLUT1-4, blockade of all glucose metabolic reactions using 2-deoxyglucose (2-DG), and fatty acid uptake via CD36 with its translocation into the mitochondria using CPT1 proteins for fatty acid oxidation (FAO) and blockade of FAO using Etomoxir (ETO). **(C)** NDS, DS, and Ph-like B-ALL CLs were exposed to DRB18 (10 μM), 2-DG (10 mM), ETO (10 μM), or DMSO vehicle for 72h and viability determined using flow cytometry. Live cells=annexin-V-/DAPI-. The % change in survival (relative to Vehicle) is shown. **(D)** Diagram depicting differential metabolic pathways glucose is used for with the genes underlined that are drug targets for these pathways. **(E)** CLs were cultured for 72h as in **C** in the presence of inhibitors for gluconeogenesis (FBPase-1 Inhibitor-1 (10 μM)), de novo Serine biosynthesis (CBR-5884 (10 μM)), pentose phosphate pathway (PPP) (6-aminonicotinamide (500 nM)), lactate metabolism (GSK-2837808A (500 nM)), hexosamine (E1276 (10 μM)), and mitochondrial pyruvate carrier (UK5099 (10 μM)). **(F)** CLs were cultured as in **E** with the addition of VEN (0.02 μM). **(G-H)** DS B-ALL CLs were transfected with GFP-Cas9 and Control-gRNA (white bars), PHGDH-AAgRNA (grey bars), or PHGDH-ABgRNA (red bars) for 60h. Viability was measured on GFP+ cells using DAPI/annexin-V staining **(G)** or after 24h treatment with VEN 0.02 μM **(H)**. Bar graphs represent average % survival ± SEM from n=3+ replicates for all treatments and CLs. One-way ANOVA with Tukey posttest was used for statistics. *p<0.05, **p<0.01, ***p<0.001, and ****p<0.0001.

## Discussion

The elevated risk of B-ALL in children with DS combined with their increased incidence of chemotherapy intolerance, treatment-related mortality, and relapse^3-5^ warrants the investigation of novel therapies. CLs have been foundational for drug testing in human cancers and surprisingly there have been no DS B-ALL CLs to study. Given this, we generated three DS B-ALL CLs that represent distinct subtypes with differing mutations, including a novel *ERG::CEBPD FLT3-*mutant. *ERG* is an ETS transcription factor on chromosome 21 frequently deleted or truncated in the *DUX4* B-ALL subset^22^. Wildtype *ERG* induced erythromegakaryoblastic leukemia^23^, whereas truncated *ERG* induced B-ALL in mice^22^. The role of *ERG* as a fusion partner with *CEBPD* warrants further investigation in DS B-ALL. Surprisingly, we found *TP53* mutations in DS-1 and DS-81 CLs, which might be explained by their relapse status^24,25^ and also warrants future studies in relapsed/refractory DS patients. Nevertheless, we found a consistent sensitivity to Venetoclax in DS regardless of mutational subsets, which included efficacy against *CRLF2-rearranged* (*BCR::ABL1-like*). These results are promising for Venetoclax therapy in DS B-ALL considering *TP53* is essential for sustaining effective BH3-mimetic responses in leukemias^26,27^ and our DS CLs had similar responses regardless of *TP53* status, which might be explained by mitochondrial dysfunction found in DS CLs that was previously reported in multiple other DS cell types^28-33^. Our DS CLs have both similarities and differences with Ph-like B-ALL CLs. Similarities include diminished proliferation and mitochondrial function. However, DS was more sensitive to Venetoclax and distinct patterns of glucose utilization were present amongst the three cohorts as NDS used glucose for glycolysis, Ph-like for PPP, while DS displayed poor use of glucose *in vitro*. In contrast, we found DS B-ALL patient samples displayed an enriched metabolic profile, especially in pathways involving glucose, fatty acids, and lipids. In addition to the metabolic differences found between DS CLs and patient samples, we observed differential expression patterns of BCL2 family members when compared to NDS.

The rationale for testing Venetoclax in Ph-like B-ALL was that anti-apoptotic *BCL2*-assocaited athanogene-1 (*BAG1*) was overexpressed in patient samples; however, expression levels of BAG1 and other BCL2 family members were unchanged amongst Ph-like and NDS B-ALL CLs^12^. We also found BCL2 family member expression levels could not explain the enhanced sensitivity to Venetoclax observed in DS B-ALL CLs. While correlating Venetoclax efficacy with BCL2 family member gene expression has yielded controversial results in other leukemia and lymphoma subtypes^34,35^, it appears DS B-ALL patients are reliant on anti-apoptotic BCL2 genes to resist apoptosis as *MCL1, BCL2L1*, and *BAG1* were elevated when compared to NDS; along with upregulation of *BAX, BAK, BAD, PMAIP1*, and *BBC3* pro-apoptotic family members. In contrast to Ph-like B-ALL where Venetoclax was only effective *in vivo* when combined with Ruxolitinib^12^, we show efficacy as a single agent in mice challenged with DS B-ALL CLs and PDXs. The efficacy of Venetoclax in other NDS B-ALL subtypes is controversial and generally ineffective as a single agent^17,36-39^, whereas others recently found Venetoclax to be effective using DS B-ALL PDXs *in vitro*^40^.

Vincristine has been a cornerstone therapy in B-ALL for decades. Here, we show synergy with Venetoclax and Vincristine both *in vitro* and *in vivo* for DS B-ALL. Interestingly, both drugs were effective *in vitro* at low nanomolar concentrations and their combination *in vivo* resulted in almost complete eradication of DS B-ALL. These results support the testing of Venetoclax in combination with Vincristine in relapsed/refractory DS B-ALL. Additionally, we found that <16% of B-ALL patient samples were in the S-G_2_-M phase of the cell cycle, suggesting that multiple cell cycle directed agents, requiring cell proliferation (Vincristine, Methotrexate, and Doxorubicin) might not be beneficial. Instead, combining Venetoclax with drugs targeting constitutively active cell signaling pathways, such as Trametinib or Ruxolitinib, appear logical and have been proposed for DS^16,41^ and Ph-like B-ALL^12,42^, respectively. Indeed, we found Venetoclax to synergize with Trametinib in DS B-ALL and suggest this over Ruxolitinib because it targets the constitutively active RAS-ERK pathway found in DS^41^, while also enhancing T-cell mediated tumor clearance^43,44^ that would be inhibited by Ruxolitinib^45-47^.

An interesting finding from this study was the altered metabolic phenotype observed in DS B-ALL patients, mainly involving lipids and glucose metabolism. We find glucose is critical for B-ALL survival, that others also found in NDS CLs^48^. It appears that glucose is critical in B-ALL to support *de novo* serine biosynthesis, that was also observed in T-ALL^49^ and other cancers^50-52^. While asparaginase therapy in B-ALL has been efficacious for decades, targeting serine metabolism might provide a therapeutic window for relapsed/refractory B-ALL patients that warrants future investigations. A weakness of this study was our limited number of B-ALL patient samples used for metabolomic analysis; however, we interrogated the transcriptomes of DS B-ALL patients (n=249)^6^ and NDS (n=1988)^15^, that showed multiple genes involved in glucose and fatty acid metabolism were upregulated in DS, supporting our mass spectrometry findings. Although some of our findings in DS B-ALL patient samples and CLs contradicted each other when compared to similar sample types from NDS, our DS CLs have similar defects in cell proliferation and mitochondrial function as commercially available Ph-like CLs used for decades, supporting their use for DS biology studies.

In summary, we have: (1) generated novel tools to study DS B-ALL by creating and characterizing three distinct CLs; (2) conducted the first mass spectrometry-based multiomics study in pediatric B-ALL that unveiled a new layer in the biology of DS as these patients exhibit an altered metabolic profile that would presumably increase chemoresistance and rates of relapse, which are hallmarks in DS; and (3) identified Venetoclax to be effective against DS B-ALL CLs, PDXs, and patient samples. Its efficacy at low nanomolar concentrations and synergy with Vincristine supports clinical testing of Venetoclax for DS patients with relapsed/refractory B-ALL.

## Notes

### Competing Interest Statement

The authors have declared no competing interest.

### Summary of Updates

This version has been updated with a revised Figure 7.

