## Supplementary material for "Metabolic vulnerabilities in Down syndrome B-cell acute lymphoblastic leukemia can be targeted using Venetoclax": S

**Patient samples, cell culture, and GFP-luciferase transduction**

Cryopreserved B-ALL patient samples and patient derived xenografts (PDXs) were thawed and cultured as previously described[1]. Briefly, cells were thawed and cultured at 1.0 x$10^6$ cells/ml in IMDM (ThermoFisher) supplemented with 20% FBS (Sigma), 1X antibiotic/antimitotic (ThermoFisher), 1X glutamax (ThermoFisher), 55-µM beta-mercaptoethanol (ThermoFisher), 1X BIT (bovine-serum albumin, insulin, and transferrin; StemCell), IL-3 (10-ng/ml; RnD Systems), and IL-7 (10-ng/ml; Biolegend). Cell lines (CLs) were cultured in IMDM (ThermoFisher) supplemented with 10% FBS (Sigma), 1X antibiotic/antimitotic (ThermoFisher), 1X glutamax (ThermoFisher), and 55 µM beta-mercaptoethanol (ThermoFisher). DS-76 and DS-81 CLs were cultured at 1.0 x$10^6$ cells/ml and media was doubled after 3 days of culture and then cells were resuspended in fresh media at 1.0 x$10^6$ cells/ml after on day 5-6. DS-1, MUTZ5, and MHH-cALL4 CLs were cultured similarly but seeded at 2.0 x$10^6$ cells/ml. NDS B-ALL CLs were seeded at approximately 3.0 x$10^5$ cells/ml and split 1:6 every 3 days. DS B-ALL CLs were generated by culturing cells at 1.0 x$10^6$ cells/ml with 50% media changes every 24-48 h for 4-6 weeks without the addition of BIT or cytokines. Cells were expanded from 3-20 x$10^6$ cells to 1-2 x$10^8$ cells before cryopreserving for future usage. All B-ALL CLs and patient samples were cultured on tissue culture treated 6-well or 10$cm^2$ plates. Due to differing cell culture techniques between NDS, DS, and Ph-like B-ALL CLs, we seeded all three cohorts of CLs at 1.0 x$10^6$ cells/ml for 24-48 h before using for seahorse, metabolism, and cell cycle experiments conducted herein. DS and NDS B-ALL CLs were lentivirus transduced with pCDH-EF1a-eFFly-eGFP as previously described[2], followed by flow cytometry sorting of $GFP^+$ cells on a FACS Aria to allow *in vivo* imaging.

**Whole-genome and RNA sequencing from B-ALL Samples**

Whole-genome sequencing (WGS) libraries were constructed using Kapa Hyperprep library preparation kit (Roche) and sequenced on NovaSeq platform (Illumina) at St. Jude, with a targeted coverage of 60X and a read length of 2 x 151. RNA was extracted from 1-10 x$10^6$ cells from both CLs and patient samples using RNeasy mini kit (Qiagen) based on the manufacturer's protocols. RNA from CLs was extracted from freshly cultured cells while frozen patient samples were used for extraction. RNA quality was determined by BioAnalyzer (Agilent) and quantified by Qubit (Life Technologies). Samples with RIN of 7 or greater and a minimum of 500 ng total RNA were used for library prep (using 3 prime kits from 10X Genomics) and sequencing. Libraries were sequenced with an Illumina HiSeq 2000 System at the UCCC Genomics Core. Samples were submitted to the Genomics and Microarray Core at the University of Anschutz Medical Campus for 2x150 bp sequencing on an Illumina NovaSeq 6000. Reads were inspected for quality using FastQC v0.11.9 and MultiQC v1.9. Reads were then quantified using Salmon v1.4.0[3] and aggregated at the gene level using tximport v1.18.0[4] in an R v4.0.2 environment. Reads were joined with unpublished HTSeq count data and normalized using the median ratio method implemented in DESeq2 v1.30.0[5]. Differential testing was then conducted by fitting a negative binomial generalized linear model to each gene and applying the Wald test to the resulting beta estimates, as is implemented by the DESeq function. Adjusted p-values were calculated using the Benjamini-Hochberg procedure following independent filtering. RNA-seq IPA analyses was conducted as previously described[6].

**Flow cytometry**

B-ALL CLs and patient samples were surfaced stained for 30-min on ice, washed in FACS-buffer (PBS + 2% FBS) and then either resuspended in FACS-buffer and analyzed on a Cytek Aurora or fixed in 1% PFA (paraformaldehyde, ThermoFisher). For intracellular analyses of CD79A, cells were fixed and permeabilized based on manufacturer's directions (Fixation/Permeabilization Kit, BD Biosciences) and stained for their expression levels and acquired on a Cytek Aurora. Cell cycle analyses using Ki67 and DAPI staining was conducted as previously described[7]. For patient sample analyses, cells were thawed from cryopreservation and immediately surfaced stained, fixed and permeabilized and stained for Ki67 expression levels, then washed and stained using DAPI and acquired on a Cytek Aurora. For patient sample chemotherapy screens, cells were harvested and surface stained on ice for 30-min using Annexin-V staining buffer (Biolegend), washed in FACS buffer and then resuspended in DAPI and analyzed on a Cytek Aurora. For cell-trace-violet (ThermoFisher) proliferation assays cells were stained based on the manufacturer's protocols and cultured at 1.0 x$10^6$ cells/ml for 6-days with cells being split 1:1 every 48-h and acquired on a Cytek Aurora. Aside from chemotherapy experiments, the eBioscience fixable viability dye eFluor780 was used (ThermoFisher). Oxidative stress was measured in B-ALL CLs using CellRox Deep Red (ThermoFisher) after culturing in complete media for 4-hours. Cells were harvested and stained as above. A list of all flow cytometry antibodies used in this study can be found below.

| **Antibodies** | **Source** | **Identifier** |
|---|---|---|
| CD34-UV395 | BD Biosciences | 745585 |
| CD34-PE | BD Biosciences | 550619 |
| CD19-BV605 | Biolegend | 302224 |
| CD19-FITC | Biolegend | 302206 |
| CD45-BV510 | Biolegend | 304036 |
| CD45-APC-Cy7 | Biolegend | 368516 |
| CRLF2-PE | Biolegend | 322806 |
| CRLF2-APC | Biolegend | 151806 |
| CRLF2-PE-Cy7 | Biolegend | 322810 |
| CD22-PerCP-Cy5.5 | Biolegend | 302516 |
| Ki67-PE | Biolegend | 350504 |
| CD79A-APC | Biolegend | 333506 |
| FC block | Biolegend | 422302 |
| CD3-PerCP-Cy5.5 | Biolegend | 300328 |
| CD66B-FITC | Biolegend | 305104 |
| IL-7Rα-BV421 | Biolegend | 351310 |
| CD14-FITC | Biolegend | 325604 |
| AnnixinV-PE | Biolegend | 640947 |
| GLUT1-AF405 | R&D Systems | FAB1418V |
| CD36-BV605 | BD Biosciences | 563518 |

**Western Blot and Immunoprecipitation (IP)**

Whole cell lysates were extracted on ice using RIPA buffer (ThermoFisher) supplemented with dithiothreitol (DTT 1 µM, ThermoFisher), Phosphatase Inhibitor Cocktail 2 and 3 (Sigma), and cOmplete Mini Proteinase Inhibitor Cocktail (Roche) for 20-min with vortexing. Lysates were quantified using BCA assays (ThermoFisher). For western blots, 30-60 µg of protein were loaded for each cell line. ImageJ was used for all Western blot analyses. IPs were conducted using Universal Magnetic Co-IP kit based on the manufacturer's protocol (Active Motif).

| Antibodies | Source | Identifier |
|---|---|---|
| P-4EBP1 | Cell Signaling | 2855 |
| 4EBP1 | Cell Signaling | 9644 |
| P-STAT5 | Cell Signaling | 9351 |
| STAT5 | Cell Signaling | 25656 |
| P-ERK-1/2 | Cell Signaling | 9101 |
| ERK-1/2 | Cell Signaling | 4695 |
| P-BCL2 | Cell Signaling | 2827 |
| BCL2 | Cell Signaling | 4223 |
| BCL2 (IP pulldown) | BD Biosciences | 51-1513GR |
| BCL-XL | Cell Signaling | 2764 |
| MCL1 | Cell Signaling | 5453 |
| BAX | Cell Signaling | 2772 |
| BAD | Cell Signaling | 9239 |
| BIM | Cell Signaling | 2933 |
| β-Actin | Cell Signaling | 4970 |
| P-S6 | Cell Signaling | 4858 |
| S6 | Cell Signaling | 2217 |
| P-AKT | Cell Signaling | 4060 |
| AKT | Cell Signaling | 4691 |
| c-Myc | Cell Signaling | 5605 |
| BAK | Cell Signaling | 12105 |
| BAG1 | Santa Cruz | SC-56003 |
| PUMA | Cell Signaling | 12450 |
| NOXA | Cell Signaling | 14766 |
| CPT1A | Cell Signaling | 12252 |
| CPT1B | Abcam | AB134988 |

**Mouse studies**

The Venetoclax studies in NSG-mice were conducted as previously described[8,9]. Briefly, 1.0 x$10^6$ NDS or DS B-ALL CLs were tail-vein injected into NSG mice. Then, on day-10 (DS-76 and DS-81) or day-14 (DS-1) mice were treated for 4-weeks (5-days/week) with Venetoclax (100mg/kg, MedChem Express)[8] or vehicle. Bioluminescence imaging (BLI) of luciferin treated mice was conducted to monitor leukemia growth. Weekly images were obtained for Venetoclax studies and images every 10-days after tail-vein injecting B-ALL cells to compare leukemia growth between NDS and DS. DS B-ALL PDXs were generated from tail-vein injection as mentioned above and used in this study at passage-2. B-ALL in peripheral blood was monitored by tail-vein cuts and flow cytometry for human CD19 expression. Once levels were >1% human-CD19$^+$ we began treatment with vehicle, Venetoclax as above, Vincristine as previous[9] (0.5mg/kg/week), or

Venetoclax+Vincristine combo for 4-weeks. Mice were sacrificed on day-31 of treatment and spleens harvested for absolute # of B-ALL cells/spleen that was determined using flow cytometry and precision counting beads (Biolegend Cat# 424902).

**Seahorse Mito-stress test and oxygen consumption rate (OCR) and extracellular acidification rate (ECAR) measurement**.

An XFe-96 Extracellular Flux Analyzer (Agilent) were used for measurement of OCR (OXPHOS) and ECAR (glycolysis). B-ALL CLs were plated in XF DMEM Seahorse medium supplemented with 10 mM glucose and 2 mM L-glutamine at 2.5 $x10^5$ cells per well by centrifuging in Seahorse plates coated with CellTak (Corning) at 200g for 2 minutes (break off) and then incubated in a non-$CO_2$ incubator for 30 minutes. The OCR and ECAR were measured in XF DMEM Seahorse medium supplemented with 10 mM glucose and 2 mM L-glutamine in response to oligomycin (6 µM, Sigma), FCCP (2 µM, Sigma), rotenone (250 nM, Sigma) plus antimycin (165 nM, Sigma) and 2-DG (100 mM, Sigma). Data was normalized via Hoechst staining (Fisher) and imaging (BioTEK Cytation).

OXPHOS measurements were calculated via Mitochondrial Stress Test (Agilent).

Glycolytic measurements were calculated as follows:

Basal Glycolysis=$[ECAR]_{basal}$-$[ECAR]_{2DG}$

Glycolytic capacity= $[ECAR]_{oligomycin}$-$[ECAR]_{2DG}$

Spare glycolytic reserve= Glycolytic Capacity-Basal Glycolysis

**Confocal imaging and analysis of mitochondrial morphology**

B-ALL cell lines (1 $x10^6$ cells) were stained using Mitospy CMX Ros (250 nM, Biolegend) for 30 min at 37°C and fixed in 2% paraformaldehyde (PFA, Sigma) for 15 min at RT, prior to nuclear staining with 4',6-diamidino-2-phenylindole, dihydrochloride ((DAPI) 300 nM, ThermoFisher) for 5 min at room temperature. B-ALL cells were mounted using Mowiol (Sigma) and imaged on a Zeiss LSM 780 Spectral inverted motorized microscope equipped with a ×63/1.4 N.A. oil objective and 561nm and Mai-Tai 2-photon (Spectra physics) laser lines. Z-stacks at 0.27 µm increments and 0.1 µm lateral resolution were captured using PM detectors in conjunction with Zen 2012 acquisition software. Mitochondrial fluorescence and morphology were quantified in Fiji (http://fiji.sc/Fiji) as Z-stacks collapsed into Maximum Intensity Projections as previously described[10].

**Metabolic flux using $^{13}C_6$ glucose tracing, metabolomics, and lipidomics**

NDS and DS B-ALL CLs were cultured for 24-h in glucose-free DMEM (ThermoFisher) supplemented with $^{13}C_6$ glucose (10 mM, Cambridge Isotope Laboratories), 5% FBS (Sigma) and 5% dialyzed FBS (Gibco) to minimize non-labeled glucose from non-dialyzed FBS, 1X antibiotic/antimitotic (ThermoFisher), 1X glutamax (ThermoFisher), and 55-µM beta-mercaptoethanol (ThermoFisher). Then, cells were harvested and washed 3X in cold PBS and 1.0 $x10^6$ cells were analyzed in triplicate from all NDS, DS, and MUTZ5 Ph-like B-ALL CLs. Metabolomics[11,12] and lipidomics[13] analyses were performed via ultra-high pressure liquid chromatography-mass spectrometry (UHPLC-MS – Vanquish and Q Exactive, ThermoFisher) as previously reported. Metabolites were extracted in ice cold methanol:acetonitrile:water (5:3:2 v/v/v) and lipids were extracted using cold methanol at a concentration of 2 $x10^6$ cells/ml. After vortexing for 30 min at 4° C, samples were centrifuged at 15,000 g for 10 min at 4° C and supernatants processed for metabolomics using a 5-min C18 gradient in positive and negative ion modes (separate runs) exactly as previously described[12]. Metabolite assignment was performed against an in-house standard library, as reported[11]. Peak areas for $^{13}C_2$ isotopologues were corrected for the natural abundance contribution from the respective $^{12}C$ peak. Stacked bar graphs of results were prepared using GraphPad Prism v 9.0.

### Chemotherapy drugs and viability screening

Venetoclax, Vincristine, Trametinib, Dexamethasone, Methotrexate, VU0661013, Etomoxir, and DRB18 were all obtained from MedChem Express; GSK2837808A (lactate metabolism), 6-aminonicotinamide (PPP), CBR-5884 (serine), FBPase-I Inhibitor-1 (gluconeogenesis), and E1276 (hexosamine) were all obtained from Selleckchem; and UK5099 (pyruvate) was obtained from Millipore. All drugs were diluted in DMSO based on the manufacturer's instructions. 2-deoxyglucose was obtained from MedChem Express and resuspended in complete IMDM cell culture media. B-ALL CLs were seeded at 3.0 x $10^5$ cells/ml and cultured for 72h before harvesting. Cells were stained with AO/PI dye (acridine orange/propidium iodide, Nexcelom) and viability was determined using a Cellaca automated cell counter (Nexcelom). % survival=viable drug/viable vehicle. All doses were tested in triplicate and average % survival calculated. Flow cytometry staining for Annexin V and DAPI were used on patient samples and PDXs with live cells=Annexin V-negative/DAPI-negative.

### Software

Heatmaps for RNA-seq and Western blot analyses were generated using Morpheus (software.broadinstitute.org/Morpheus/), mass spectrometry metabolism heatmaps were generated using Metabolon 6.0. Statistical significance was determined using GraphPad Prism. Samples were tested for normality before statistical analysis, if normal distribution was found then two-tailed T-tests were conducted, if normal distribution was not found then Mann-Whitney U or Wilcoxon rank-sum tests were used. FlowJo was used for all flow cytometry analyses. Ingenuity Pathway Analysis (IPA, Qiagen) was used for RNA-Seq canonical pathway analyses. Synergy calculations in for CLs were conducted using synergyfinder.fimm.fi. Image-J was used for western blot quantification.

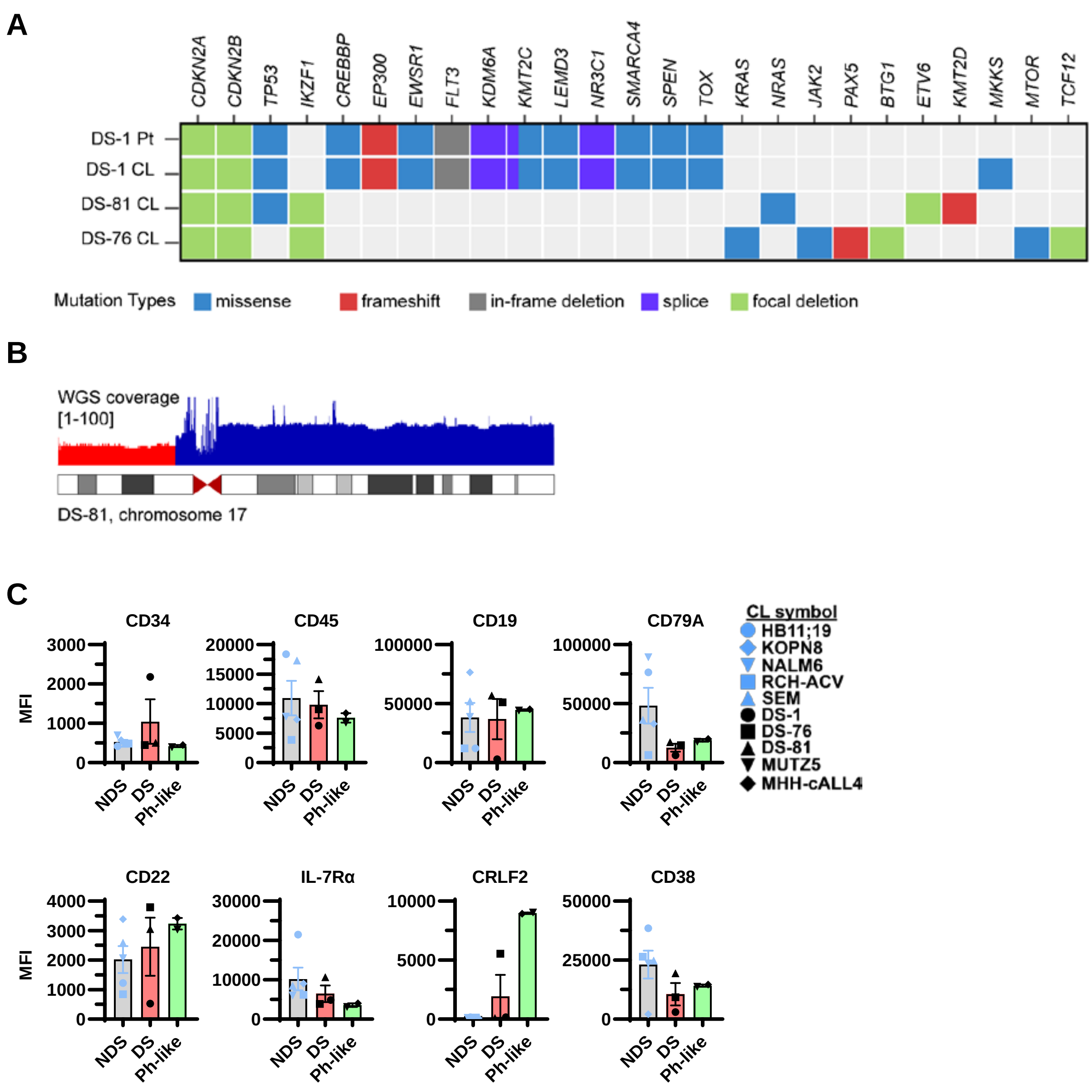


**Supplemental Figure 1. Supporting information for Figure 1. (A)** Genomic alterations of DS-1 patient sample (Pt) and the three cell lines (CLs). **(B)** Whole-genome sequencing of DS-81 CL and deletion of chromosome 17. **(C)** Flow cytometry analyses of CD34, CD45, CD19, CD79A, CD22, CRLF2, IL-7Rα, and CD38 expression levels in NDS, DS, and Ph-like B-ALL cell lines. MFI=mean fluorescence intensity. All cell lines representative of n=3+ experiments with bar graphs representing average ± SEM. Statistical significance was determined using a two-tailed T-test.

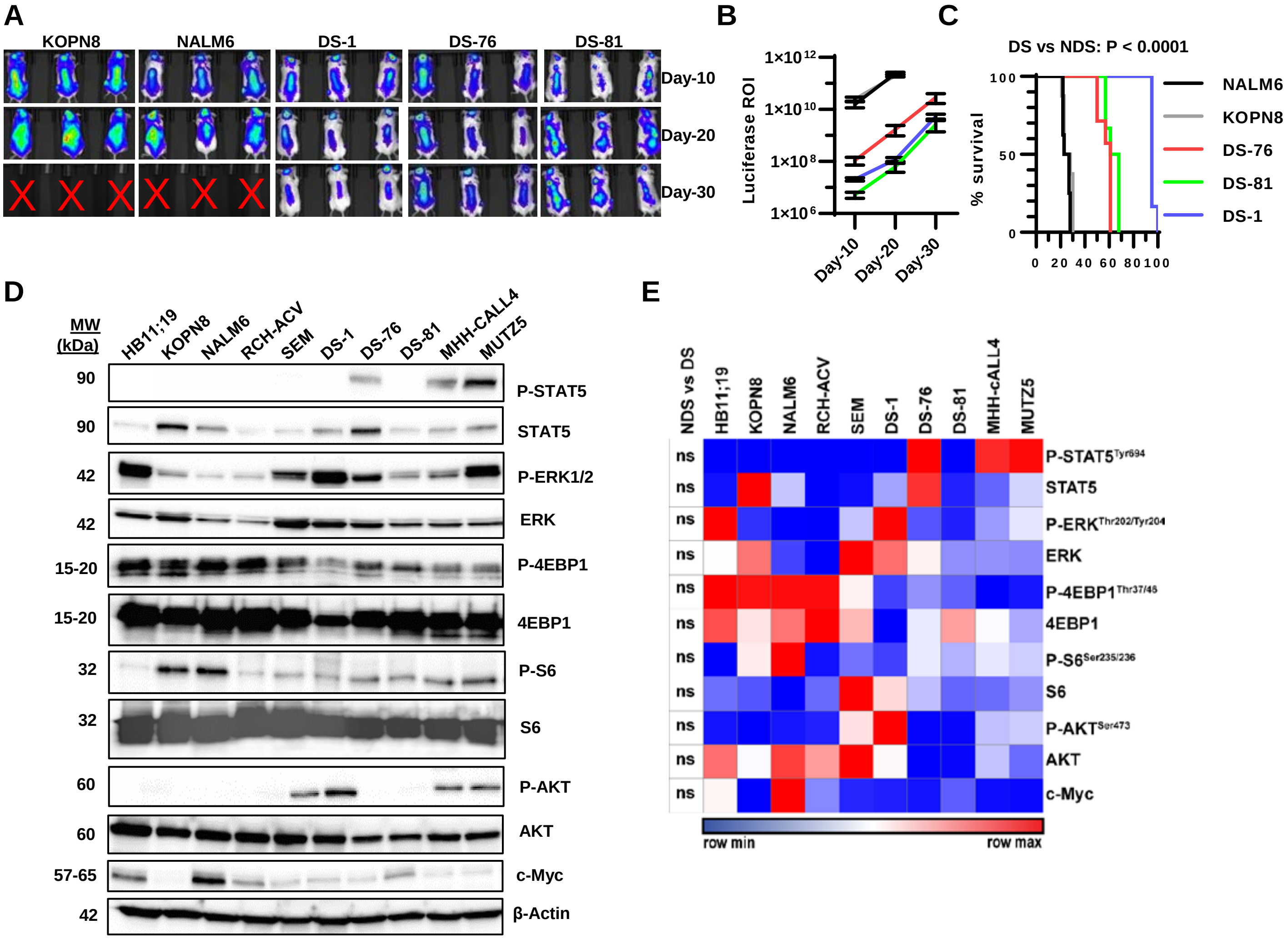


F

| Cell line | Mutation |
|---|---|
| HB11;19 | $TP53^{T125M}/TP53^{R248W}/CDKN2A^{P94L}$ |
| KOPN8 | $MLL\text{-}r/KRAS^{G12D}/TP53^{R248Q}$ |
| NALM6 | $NRAS^{A146T}/DUX4\text{-}IGH$ |
| RCH-ACV | $NSD2^{E1099K}$ |
| SEM | $MLL\text{-}r/CDKN2A^{H83Y}/TP53^{R248Q}$ |
| MHH-cALL4 | $JAK2^{I682F}/IGH\text{-}CRLF2$ |
| MUTZ5 | $JAK2^{R683G}/IGH\text{-}CRLF2$ |

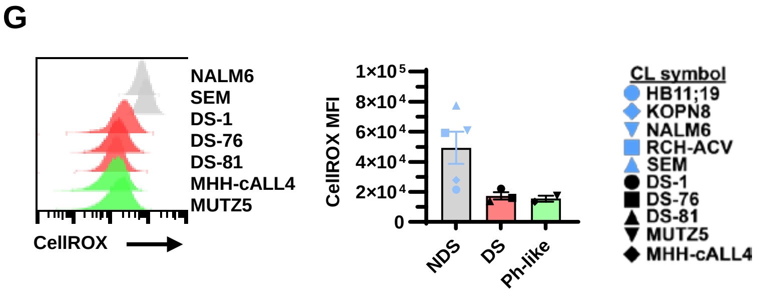


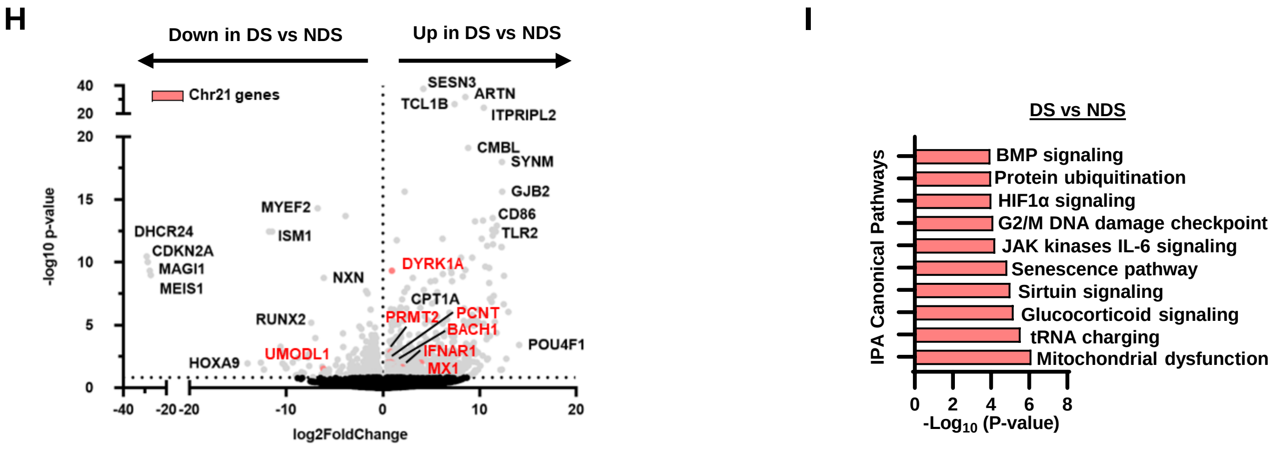

**Supplemental Figure 2. Supporting information for Figure 2. (A)** Representative bioluminescent imaging (BLI; flux measured in photons per second) of NSG mice after 10-, 20-, and 30-days of injection with 1 x $10^6$ NDS or DS B-ALL cell lines expressing GFP-Luciferase lentivirus. **(B)** Summary of luciferase region of interest (ROI) levels after 10-, 20-, and 30-days from **A**. Red X denotes dead mice. **(C)** Kaplan-Meyer survival curve from **A-B** comparing NDS and DS B-ALL cells. An n=6+ mice were used for all NDS and DS B-ALL cell lines injected into NSG mice. Log rank statistics were used to test survival differences. **(D)** Western blot analyses of phospho- (P) STAT5, ERK1/2, 4EBP1, S6, AKT and their total expression levels along with c-Myc in NDS, DS, and Ph-like B-ALL cell lines. **(E)** Quantification of western blot expression levels using Image-J from **D**. Expression levels were normalized to β-Actin from an n=2+ replicates for each protein and cell line analyzed. **(F)** Mutational analyses of NDS and Ph-like B-ALL CLs obtained from cellosaurus.org. **(G)** Representative flow cytometry histograms for CellROX analyses of oxidative stress and corresponding graph summarizing differences between NDS, DS, and Ph-like B-ALL CLs. All CLs represent an n=3+ experiments after culturing for 4h in presence of CellROX. Bar graphs=average ± SD. **(H)** Volcano plot from RNA-Seq analyses conducted on DS B-ALL CLs (DS-1, DS-76, and DS-81) compared to NDS B-ALL CLs (KOPN8, NALM6, and SEM). Significantly altered genes are shown in grey with Chr21 genes shown in red, and non-significant genes shown in black. Wald test and Adj. p-values used for statistics. **(I)** RNA-Seq analyses from **D** was subjected to IPA and the top-10 canonical pathways affected in DS B-ALL CLs compared to NDS B-ALL CLs are displayed. Statistical significance was determined using a two-tailed T-test in **E** and **G**.

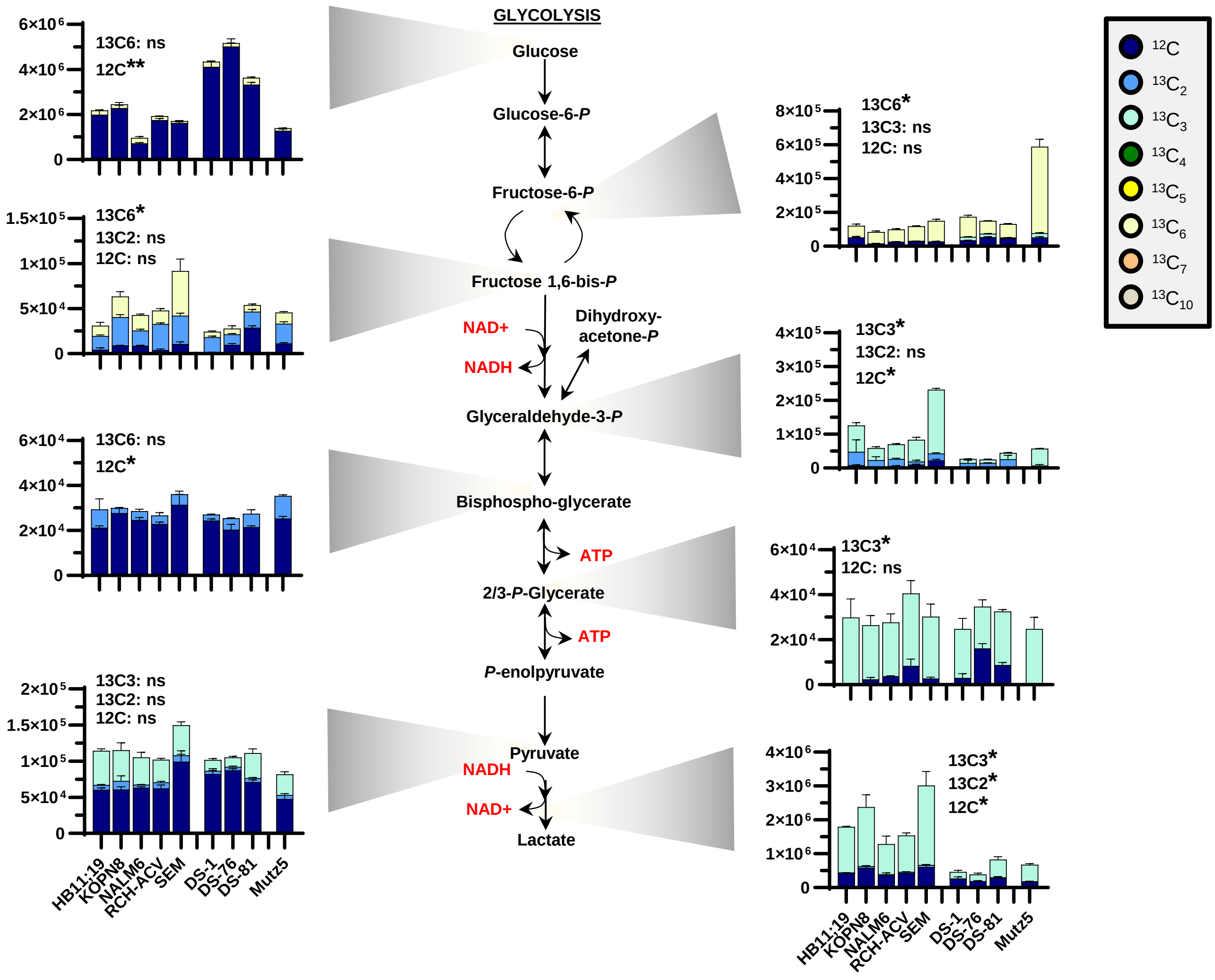


**Supplemental Figure 3. Glucose tracing in the glycolysis pathway in NDS, DS, and MUTZ5 Ph-like B-ALL cell lines from Figure-3.** The glycolysis metabolic pathway is displayed for the 24h glucose tracing mass spectrometry experiments conducted using NDS, DS, and MUTZ5 B-ALL cell lines. Both the total- and labeled metabolites in the glycolytic pathway are shown in color coated bar graphs for the NDS and DS B-ALL cell lines. Figure legend represents non-labeled and labeled-glucose found from mass-spectrometry analyses. Arbitrary units of measurement used for all metabolite readouts. Bar graphs represent average ± SD from triplicate experiments. Statistical significance was determined using a two-tailed T-test comparing NDS metabolites versus DS B-ALL metabolites. *p<0.05 and ns=not significant.

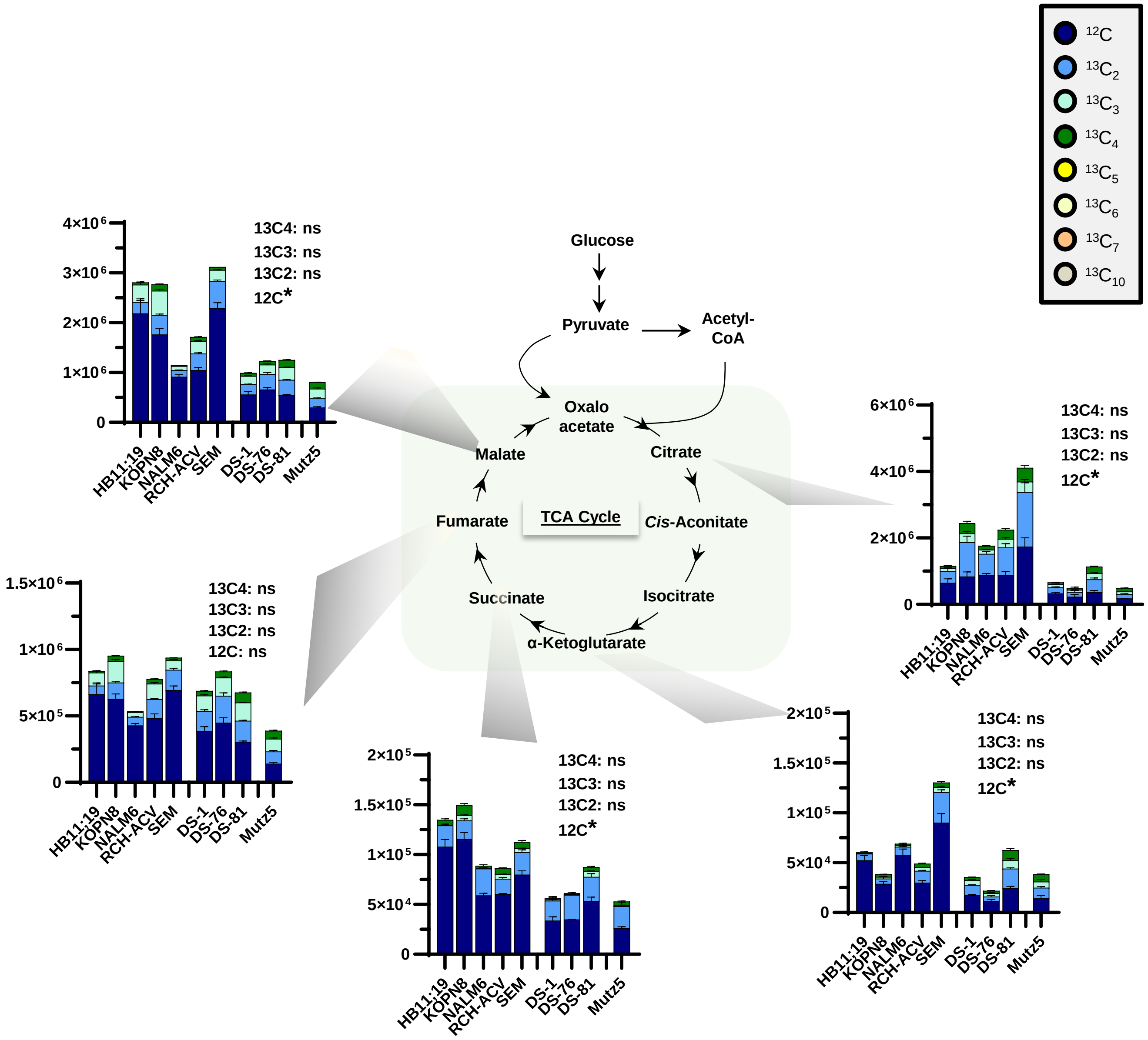


**Supplemental Figure 4. Glucose tracing in the TCA cycle in NDS, DS and MUTZ5 Ph-like B-ALL cell lines from Figure-3.** Overview of glucose metabolism into the TCA cycle in NDS, DS, and MUTZ5 B-ALL cell lines. Data in all plots are presented as peak areas (arbitrary units). Both the total- and labeled metabolites in the glycolytic pathway are shown in color coated bar graphs. Figure legend represents non-labeled and labeled-glucose found from mass-spectrometry analyses. Arbitrary units of measurement used for all metabolite readouts. Bar graphs represent average ± SD from triplicate experiments. Statistical significance was determined using a two-tailed T-test comparing NDS metabolites versus DS B-ALL metabolites. *p<0.05 and ns=not significant.

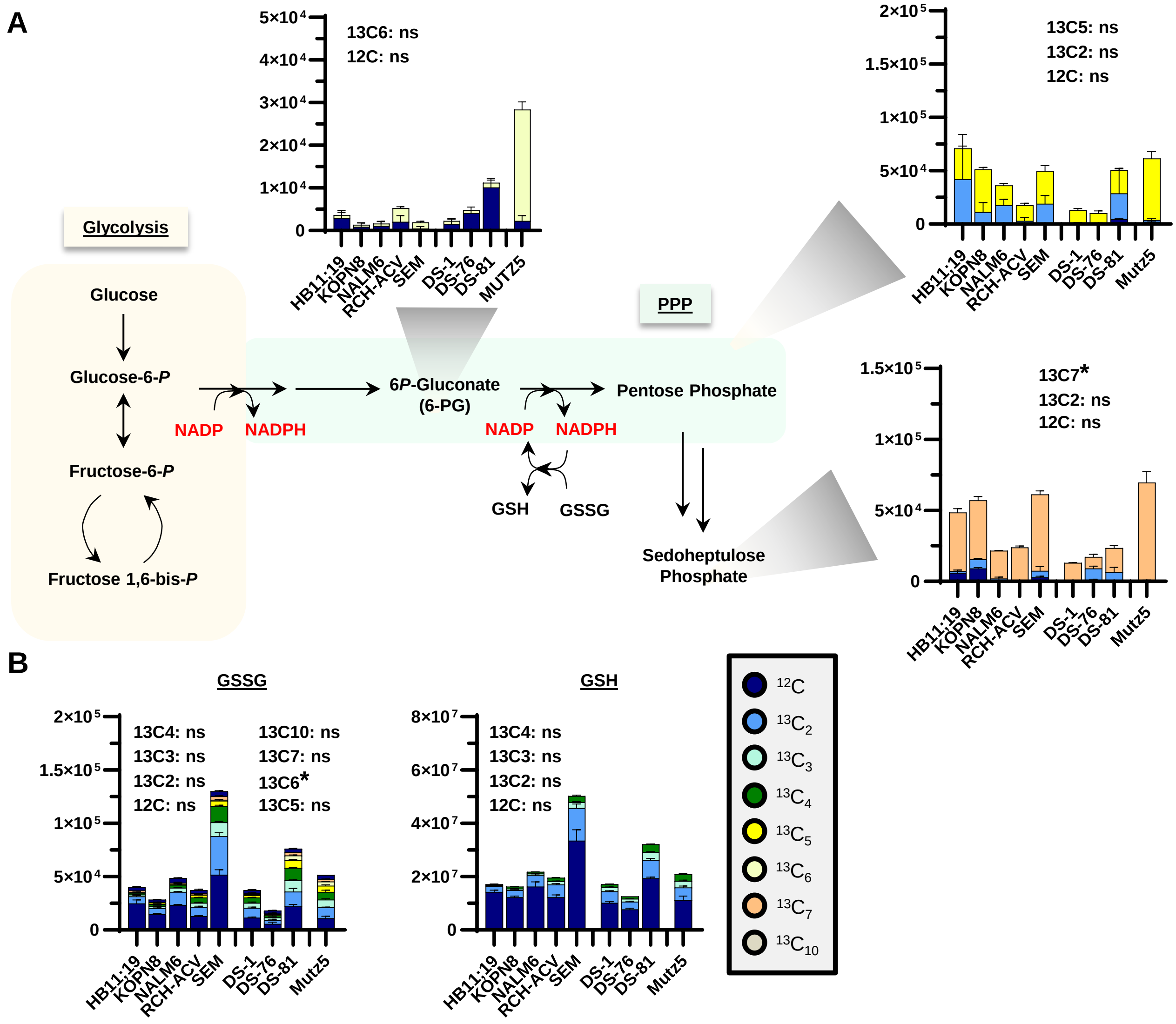


**Supplemental Figure 5. Glucose tracing in the pentose phosphate pathway and glutathione pathway in NDS, DS, and MUTZ5 Ph-like B-ALL cell lines from Figure-3. (A)** Overview of the early steps in glycolysis and how glucose can be utilized in the pentose phosphate pathway (PPP) in NDS, DS, and MUTZ5 B-ALL cell lines. **(B)** Overview of glucose tracing experiments with levels of reduced (GSH) and oxidized (GSSG) glutathione. Data in all plots are presented as peak areas (arbitrary units). Both the total- and labeled metabolites in the glycolytic pathway are shown in color coated bar graphs. Figure legend represents non-labeled and labeled-glucose found from mass-spectrometry analyses. Arbitrary units of measurement used for all metabolite readouts. Bar graphs represent average ± SD from triplicate experiments. Statistical significance was determined using a two-tailed T-test comparing NDS metabolites versus DS B-ALL metabolites. *p<0.05 and ns=not significant.

| B-ALL | Diagnosis | Cytogenetics | CRLF2-r | Age/Gender | BM/PB |
|---|---|---|---|---|---|
| NDS-1 | Primary | 46 | Yes | 3/M | BM |
| NDS-2 | Primary | 46, CDKN2A-del | No | 10/F | BM |
| NDS-3 | Primary | 46 | No | 16/M | BM |
| NDS-4 | Primary | 46 | No | 11/M | BM |
| NDS-5 | Primary | 46 | Yes | 18/M | BM |
| NDS-6 | Primary | 46, CDKN2A/B-del | No | 16/F | BM |
| NDS-76 | Relapse | 46, BCR-ABL | No | 16/F | BM |
| NDS-81 | Primary | 46, ETV6-RUNX1 | No | 6/M | BM |
| NDS-9 | Primary | 46 | Yes | 5/M | BM |
| NDS-10 | Primary | 46, CDKN2A-del | Yes | 8/F | BM |
| NDS-11 | Primary | 46 | No | unknown | BM |
| NDS-12 | Primary | 46, BCR-ABL CDKN2A-del | No | unknown | BM |
| NDS-13 | Primary | 46, ETV6-RUNX1 | No | unknown | BM |
| NDS-14 | Primary | 46, TCF3-PBX1 | No | unknown | BM |
| NDS-16 | Primary | 46, CDKN2A-del | No | unknown | BM |
| DS-1 | Relapse, CAR-T 2X | 47 | No | ?/M | BM |
| DS-2 | Primary | 47 | Yes | unknown | BM |
| DS-3 | Primary | 47 | Yes | 7/M | BM |
| DS-4 | Relapse | 47 | No | 16/M | BM |
| DS-5 | Primary | 47 | Yes | 3/M | BM |
| DS-76 | Primary | 47, CDKN2A-del | Yes | 19/M | PB |
| DS-81 | Relapse | 48 (T8 & T21) CDKN2A-del | No | 11/M | BM |
| DS-9 | Primary | 47 | No | 6/M | BM |
| DS-10 | Primary | 47, ETV6-RUNX1 | No | 6/M | BM |
| DS-11 | Primary | 47 | Yes | 4/M | BM |
| DS-12 | Primary | 47 | YES | 1/M | BM |

**Supplemental Figure 6. Characteristics of NDS and DS B-ALL patient samples used for this study.** CAR-T=chimeric antigen receptor T cell therapy (CD19- and CD19/CD22-targeted); M=male; F=female; BM=bone marrow; PB=peripheral blood. Age is given in years.

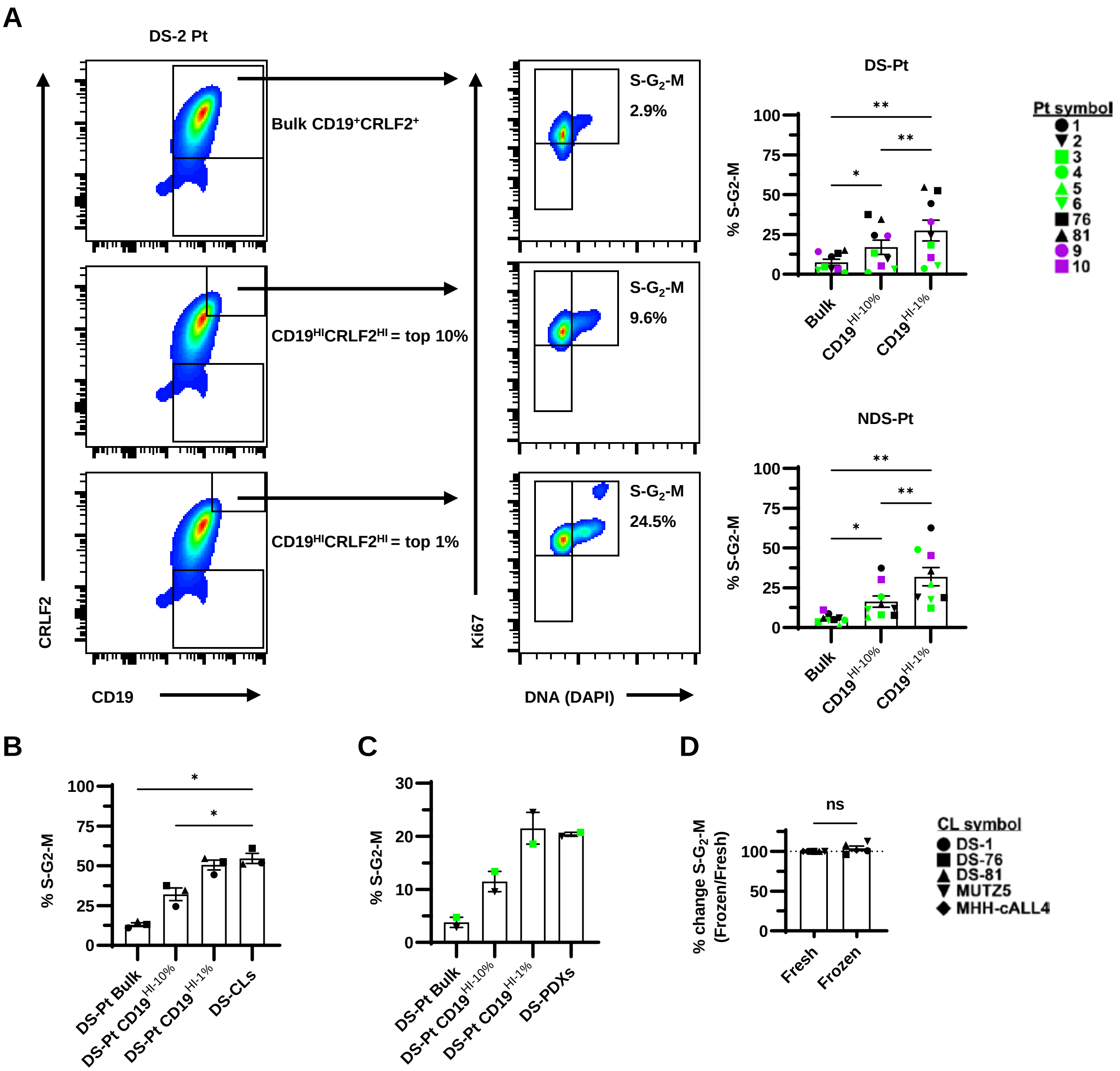


**Supplemental Figure 7. Cell cycle comparisons between patient samples, cell lines and PDXs; along with comparisons between fresh and frozen. (A)** Representative flow cytometry plot for CD19 and CRLF2 gating in DS-2 patient sample for bulk $CD19^{+}CRLF2^{+}$, $CD19^{HI}CRLF2^{HI}$ = top 10%, and $CD19^{HI}CRLF2^{HI}$ = top 1% cells and the cell cycle differences amongst these populations using Ki67 and DNA (DAPI) staining as in Figure 4A. Corresponding graphs summarizing S-$G_2$-M phase of the cell cycle differences in NDS and DS B-ALL patients comparing BULK $CD19^{+}$ cells (Bulk), $CD19^{HI\text{-}10\%}$, and $CD19^{HI\text{-}1\%}$ cells. If patients were $CRLF2^{+}$ they were denoted $CD19^{HI}$ also because $CRLF2^{HI}$ = $CD19^{HI}$. DS-1 cells were gated on $CD34^{HI}$. **(B)** Summary of S-$G_2$-M phase of the cell cycle in DS B-ALL patients 1, 76, and 81 from **A** in comparison to their matching Bulk DS B-ALL cell line levels (DS-CLs) from Figure 2B. **(C)** Summary of S-$G_2$-M phase of the cell cycle in DS B-ALL patients 2 and 3 from **A** in comparison with their matching Bulk DS PDXs levels (DS-PDXs). Results representative of an n=3+ experiments using DS PDXs at passage-2. **(D)** Summary of % change in S-$G_2$-M phase of the cell cycle in DS and Ph-like B-ALL cell lines freshly cultured or frozen before flow cytometry staining (Frozen/Fresh % change shown). Results representative of duplicate cell cycle stains from each cell line and condition. Statistical significance was determined using repeated-measured one-way ANOVA with Tukey posttest **A-C** or Two-tailed t-test **D**. *$p<0.05$ and **=$P<0.01$.

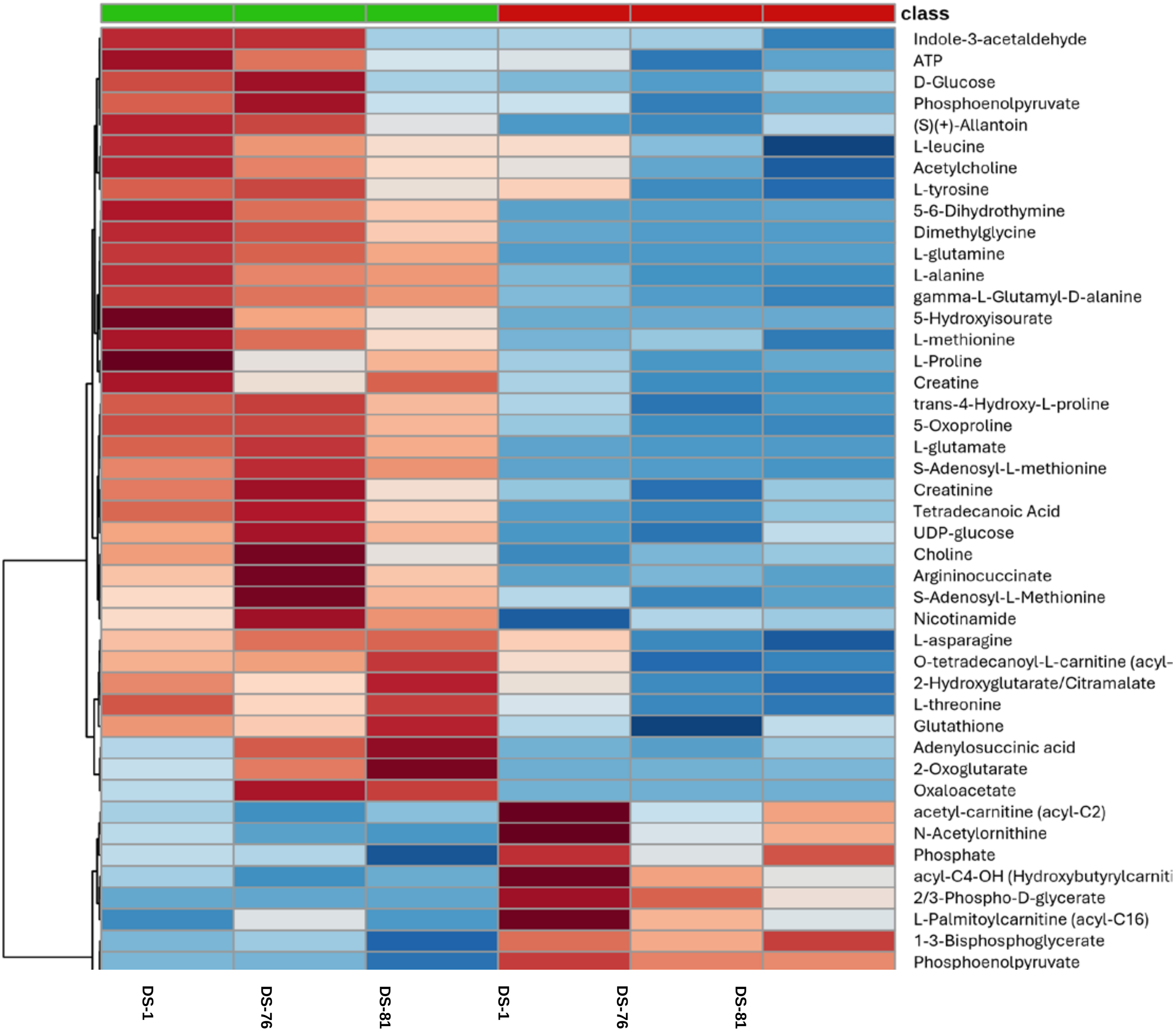


**Supplemental Figure 8. Summary of metabolites from mass spectrometry experiments comparing DS B-ALL cell lines with their matching patient samples from Figure 4B.** Cell lines are shown in green and patient samples shown in red. The top-50 metabolites based on two-tailed t-test are shown.

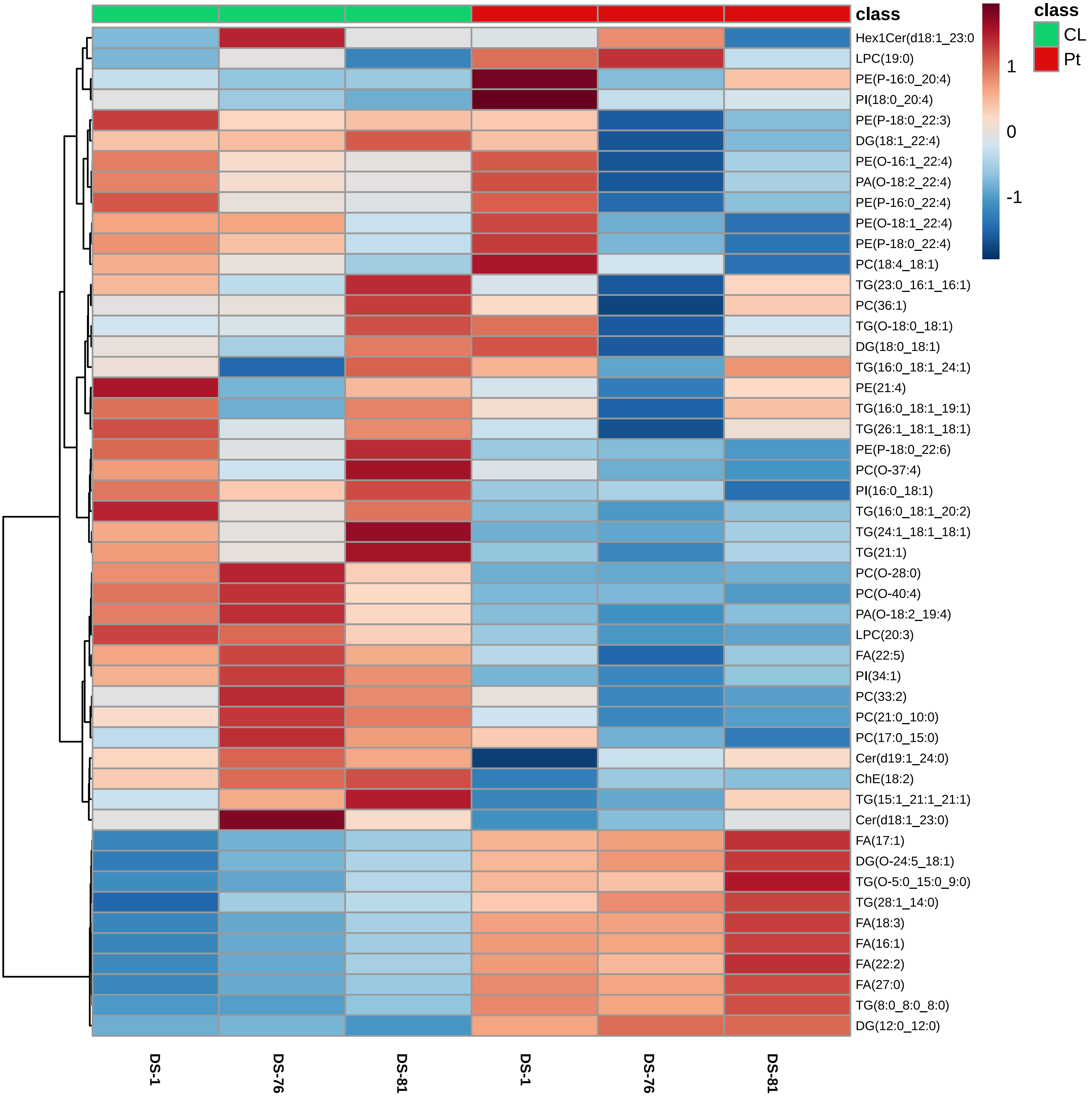


**Supplemental Figure 9. Summary of significantly altered lipids from mass spectrometry experiments comparing DS B-ALL cell lines with their matching patient samples from Figure 4C.** Cell lines are shown in green and patient samples shown in red. PE=phosphatidylethanolamine; PI=phosphatidylinositol; DG=diacylglycerol; PA=phosphatidic; TG=triglyceride; LPC=lysophosphatidylcholine; FA=fatty acid; Cer=ceramide; ChE=cholesteryl ester. Two-tailed t-test used for statistics.

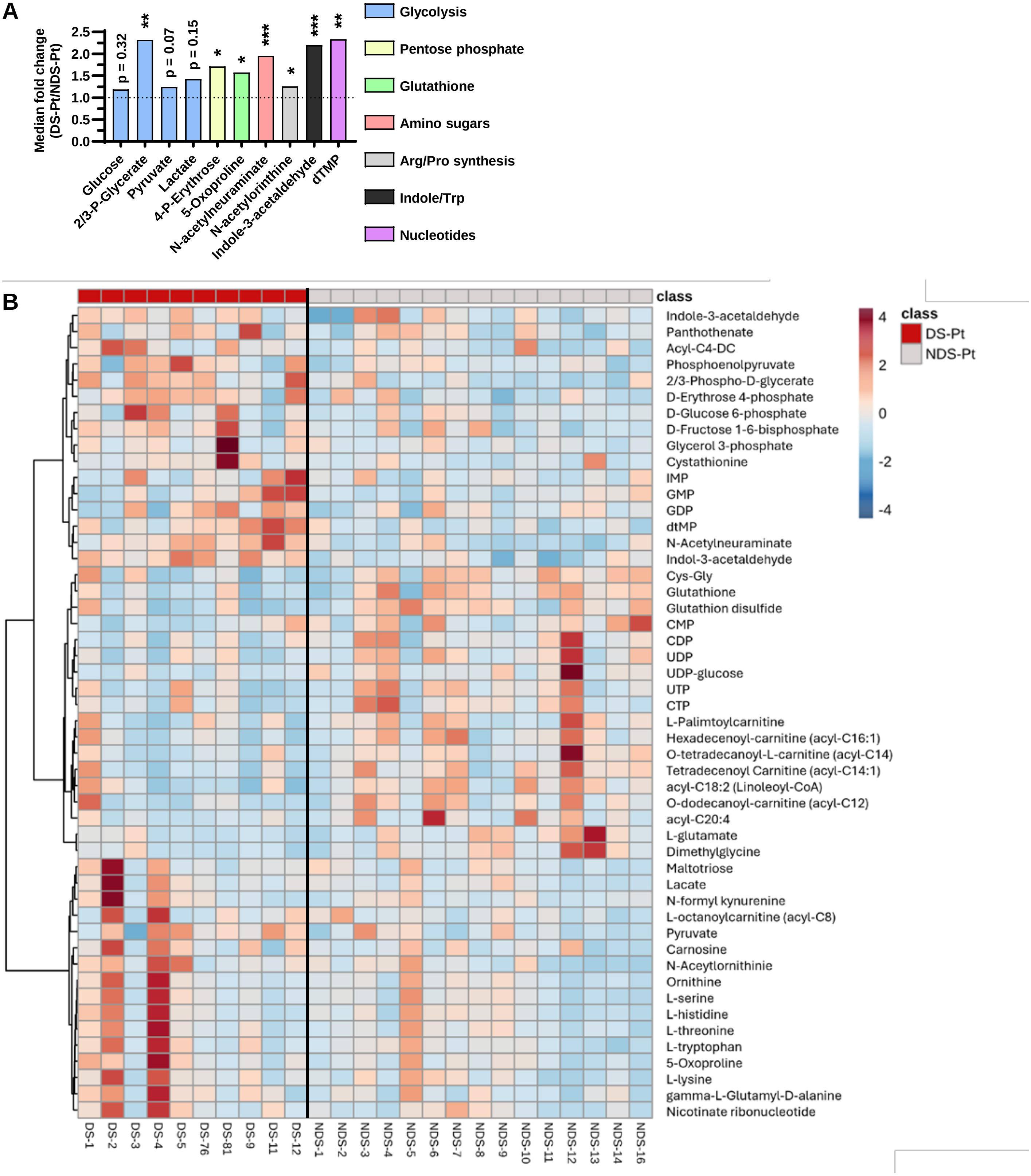


**Supplemental Figure 10. Summary of metabolite differences between DS and NDS B-ALL patients from Figure 4D. (A)** Summary of significantly altered metabolites from Figure 4D. *p<0.05, **p<0.01, and ***p<0.001. **(B)** Heatmap of metabolite differences between DS and NDS B-ALL patients from Figure 4D. DS B-ALL patients are shown in red and NDS B-ALL patients are shown in grey. The top-50 metabolites based on two-tailed t-test are shown.

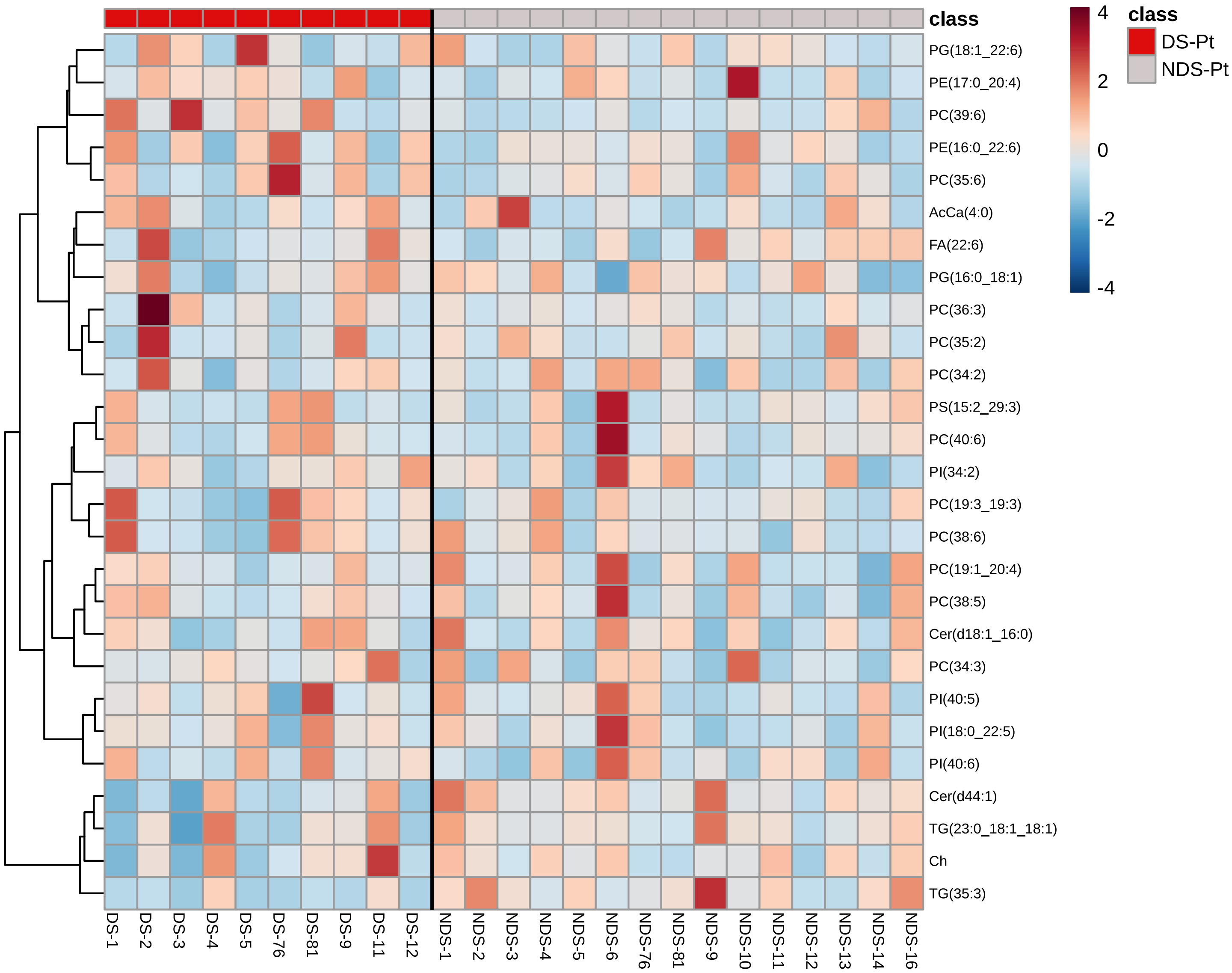


**Supplemental Figure 11. Summary of significantly altered lipids from mass spectrometry experiments comparing DS and NDS B-ALL patients from Figure 4E.** DS B-ALL patients are shown in red and NDS B-ALL patients are shown in grey. Two-tailed t-test used for statistics.

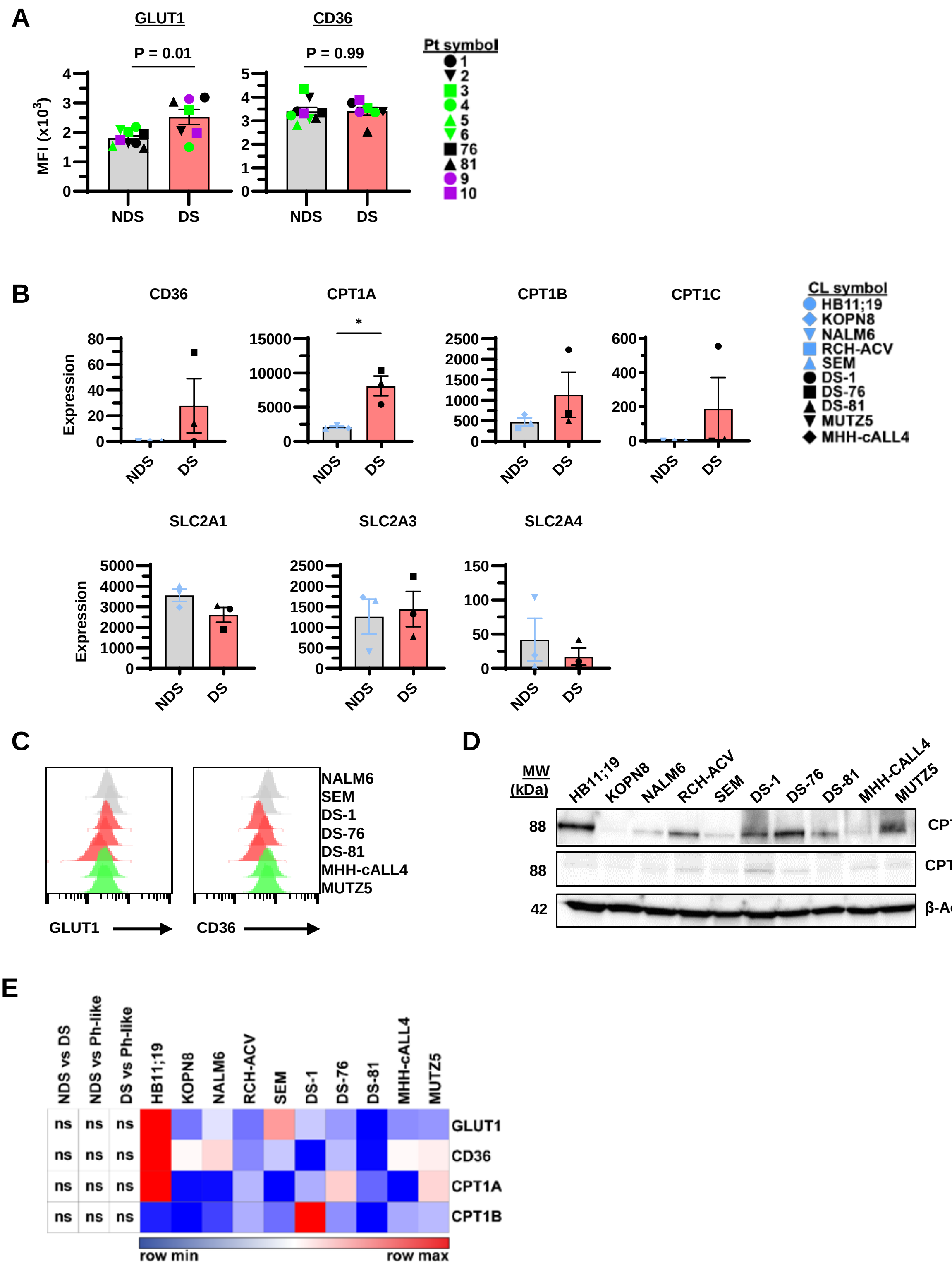


**Supplemental Figure 12. Supporting data for Figure 4. (A)** Flow cytometry analyses of GLUT1 and CD36 expression levels in DS and NDS B-ALL patient samples stained as described in Figure 4. **(B)** RNA-Seq analyses of CD36, CPT1A-C, and SLC2A1-3 expression levels in DS B-ALL cell lines (CLs) DS-1, DS-76, and DS-81 compared to NDS B-ALL CLs KOPN8, NALM6, and SEM. Bar graphs represent average ± SEM. Wald test and Adj. p-values used for statistics. **(C)** Representative flow cytometry histograms for GLUT1 (SLC2A1) and CD36 expression in NDS, DS, and Ph-like CLs. Cells were gated on live singlets. Results representative of n=3+ experiments for all CLs. **(D)** Western blot analyses of CPT1A and CPT1B expression levels. β-Actin was used as a loading control and results representative of n=2+ experiments. **(E)** Heatmap depicting changes in GLUT1 and CD36 expression levels from **C** and CPT1A and CPT1B expression levels normalized to β-Actin from **D**. Statistical significance was determined using a two-tailed t-test used for statistics. *p<0.05.

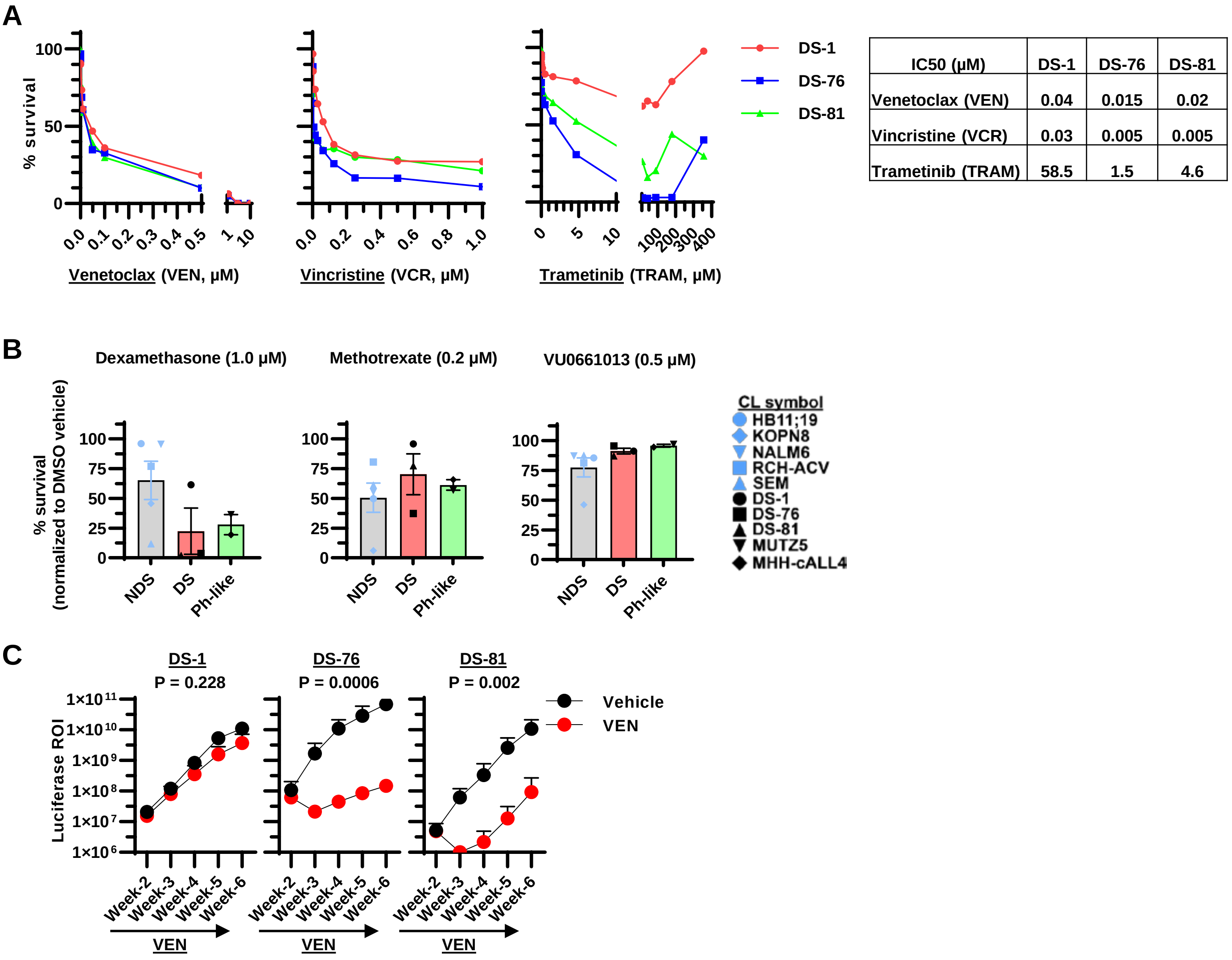


| IC50 (μM) | DS-1 | DS-76 | DS-81 |
|---|---|---|---|
| Venetoclax (VEN) | 0.04 | 0.015 | 0.02 |
| Vincristine (VCR) | 0.03 | 0.005 | 0.005 |
| Trametinib (TRAM) | 58.5 | 1.5 | 4.6 |

**Supplemental Figure 13. Supporting chemotherapy for Figure 5. (A)** Dose response curves for % survival after 72h treatment with Venetoclax (VEN), Vincristine (VCR), and Trametinib (TRAM) in DS-1, DS-76, and DS-81 B-ALL cell lines (CLs); along with their $IC_{50}$ values. CLs were stained with AO/PI dye (acridine orange/propidium iodide) and viability was determined using a Cellaca automated cell counter. % survival=viable drug/viable vehicle. All doses were tested in triplicate and average % survival shown. All drug doses are given in μM**. (B)** Graphs for % survival in response to Dexamethasone (1 μM), Methotrexate (0.2 μM), and the MCL1 inhibitor VU0661013 (0.5 μM) in NDS, DS, and Ph-like B-ALL CLs treated for 72h as in Figure-4. Cells were stained and analyzed as in **A. (C)** Summary of weekly luciferase ROI values from BLI in Vehicle or VEN treated DS-1, DS-76, or DS-81 NSG mice from Figure 5B-C. All bar graphs in **B** and time points in **C** are representative of average ± SEM. Statistical significance was determined using a two-tailed t-test in **B** and Wilcoxon rank sum test in **C**. *p<0.05.

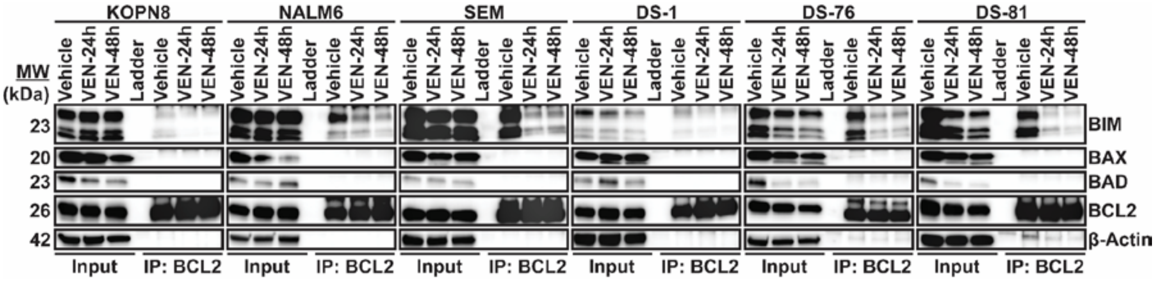


**Supplemental Figure 14. Supporting BCL2 immunoprecipitation experiments for Figure 6.** DS and NDS B-ALL cell lines were cultured at 2 x$10^6$ cells/ml and treated with Venetoclax (VEN, 50-μM) for 24- and 48h or vehicle control. Cells were harvested and whole cell lysates extracted. Then, cell lysates were subjected to immunoprecipitation (IP) with a BCL2 antibody and Western blot analyses conducted for BCL2 protein interactions with BIM, BAX, or BAD. Confirmation of the IP was conducted using western blot for BCL2 expression and total expression levels of BIM, BAX, BAD, and BCL2 were conducted in parallel experiments (Input). β-Actin was used as a loading control.

**A**

| | IC$_{50}$ values 2-DG (mM) | DRB18 (uM) |
|---|---|---|
| HB11;19 | 4.263 | 4.053 |
| KOPN8 | 3.747 | 5.781 |
| NALM6 | 1.828 | 2.522 |
| RCH-ACV | 2.196 | 4.55 |
| SEM | 5.209 | 3.754 |
| DS-1 | 1.249 | 4.423 |
| DS-76 | 5.234 | 6.1 |
| DS-81 | 4.396 | 3.21 |
| MHH-cALL4 | 4.295 | 4.568 |
| MUTZ5 | 6.579 | 6.361 |

**B**

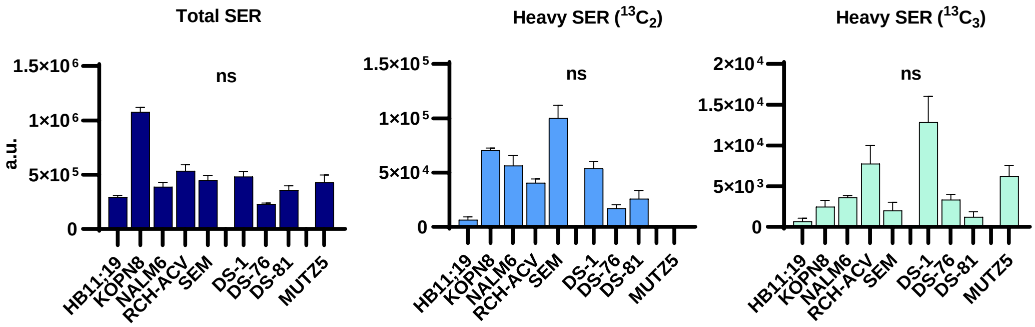


**C**

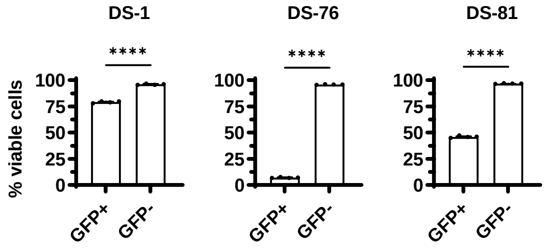


**Supplemental Figure 15. Supporting glucose and serine metabolism data for Figure 7. (A)** The IC$_{50}$ values from NDS, DS, and Ph-like B-ALL CLs treated with increasing amounts of 2-deoxyglucose (2-DG) or the GLUT inhibitor DRB18. Viability was determined using flow cytometry staining for DAPI and annexin-V (viable cells=DAPI$^-$/annexin-V$^-$). All doses were tested in triplicate. **(B)** Results from glucose tracing for total serine (SER) and labeled (heavy) SER $^{13}C_2$ and $^{13}C_3$ in B-ALL CLs. Results representative of triplicate experiments. a.u.=arbitrary units of measurement. **(C)** Results of flow cytometry viability staining in DS B-ALL CLs after 60h of CRISPR Cas9-GFP and Control-gRNA transfection. Cells were stained as in **A** and gated on GFP$^+$ and GFP$^-$ cells to show the impact of CRISPR transfection on survival. Statistical significance was determined using a two-tailed T-test. ns=not significant.
